# TRF2 mediates force-resistant bridging of distal telomeric DNA repeats

**DOI:** 10.64898/2026.10.09.757630

**Authors:** Matthew D. Newton, Thomas E. Catley, Sylvia Whittle, Lorenzo Galanti, Laura Wiggins, Elizabeth P. Holmes, Dima Daccache, Roopesh Anand, Ellen G. Allwood, Mark Greensill, Sandra Segura-Bayona, Ondrej Belan, Max C. Gamill, Sarah Maslen, Mark Skehel, Alice L. B. Pyne, Simon J. Boulton

## Abstract

Shelterin binds telomeres, regulates telomere length, and protects chromosome ends from mis-repair, which is essential for genome stability. Telomere end-protection uniquely depends on the TRF2 subunit, which assembles telomeric DNA into a protective T-loop structure *in vivo*, yet the biochemical mechanism underlying this process remains unresolved. Using single-molecule optical tweezers and atomic force microscopy, we show that the human shelterin complex tethers distal telomeric DNA repeats, looping the intervening DNA. These bridges withstand tens of piconewtons of applied load, rupture at a force that scales with telomeric repeat number and re-form on relaxation. Structure-function analysis demonstrates that this force-resistant tethering is TRF2-dependent under our conditions and depends on multiple interfaces, with the TRFH domain making a major but non-exclusive contribution: introducing the TRF2 TRFH domain into TRF1 is sufficient to confer tethering, whereas its deletion from TRF2 reduces but does not abolish activity. We propose telomeric dsDNA tethering by TRF2 as a candidate first step in T-loop formation, preceding invasion of the telomeric 3’ ssDNA overhang.

## Introduction

Human telomeres are composed of ∼10-15Kb of double-stranded (TTAGGG)_n_ repeats, ending in a G-rich single-stranded overhang, which is bound by the shelterin complex^1^. Shelterin consists of six proteins: TRF1, TRF2, TIN2, TPP1, POT1 and RAP1 (Figure 1A). TRF1 and TRF2 bind to telomeric dsDNA repeats^2–4^, whereas POT1 binds the 3’ ssDNA overhang^5,6^. These single and double-stranded DNA binding proteins are connected by RAP1, TIN2 and TPP1 to form the full shelterin complex^7–9^, with each subunit performing specialised functions that facilitate telomere replication, telomere length maintenance and telomere protection.

**Figure 1.**
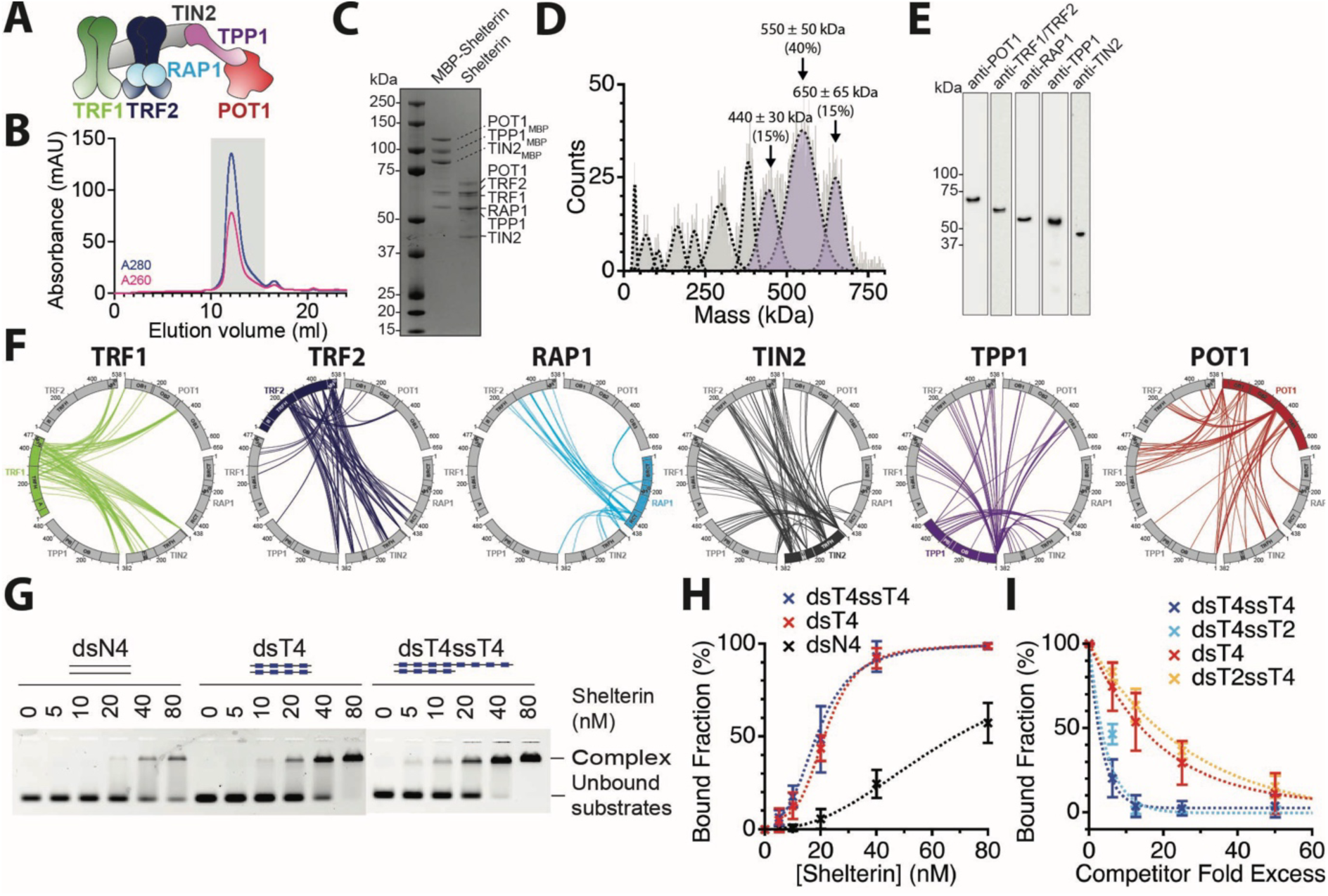
Reconstituted shelterin complexes contain both TRF1 and TRF2 and are active in telomeric dsDNA binding. (A) Schematic of the shelterin complex. (B) Chromatogram for size exclusion chromatography shows a single peak. (C) SDS-PAGE gel of pooled fractions before and after MBP cleavage, showing presence of all six shelterin proteins. (D) Mass photometry of purified shelterin complex. (E) Western blot of purified shelterin complex with anti-bodies against each of the six shelterin proteins. (F) Crosslinking mass spectrometry analysis of purified shelterin complex with 100X DSBU treatment. n = 1 (G) Agarose gel EMSA of purified shelterin binding to different telomeric and non-telomeric DNA substrates. dsN4 = 4 x double-stranded scrambled telomeric repeats, dsT4 = 4 x double-stranded telomeric repeats, dsTT4 = 4 x double-stranded telomeric repeats with 4 x single-stranded telomeric repeat overhang. (H) EMSA quantification of reconstituted shelterin binding to different telomeric and non-telomeric DNA substrates. Error bars are standard deviation (n = 3), fit with Hill fit. (I) Competition EMSA quantification of reconstituted shelterin bound to four double-stranded telomeric repeats competed with different telomeric DNA substrates. Error bars are standard deviation (n = 3), fit with exponential fit.

Through binding to telomeres, shelterin solves two major challenges at chromosome ends: the end-replication problem and the end-protection problem^1,10^. During lagging strand DNA synthesis, the replication machinery is unable to replicate to the end of linear chromosomes, which results in the loss of telomeric sequences from chromosome ends during each cell cycle^10^. Shelterin solves the end-replication problem by recruiting telomerase, a reverse transcriptase that adds telomeric sequences to maintain chromosome ends and prevent progressive shortening^9,11^. Shelterin also prevents recognition of chromosome ends as DNA damage and activation of the DNA damage response (DDR), which would otherwise lead to chromosome end-to-end fusions and cell death. Shelterin solves the end-protection problem by directly inhibiting DDR activation at telomeres^11^ and remodelling telomeres into a T-loop structure, which forms via invasion of the 3’ ssDNA overhang into internal telomeric repeats, preventing the telomere end from being recognized as DNA damage^12–16^.

Super-resolution imaging has shown that T-loops exist at the majority of chromosome ends, and conditional genetic deletion has established that, of the six shelterin subunits, only TRF2 is essential for telomeres to adopt a T-loop structure *in vivo*^17,18^. Cells inactivated for TRF1, RAP1 or POT1a/b assemble T-loops at a similar frequency to that seen in wild-type cells^17^. Furthermore, in the absence of the other shelterin proteins, TRF2 alone is sufficient for T-loop formation, and this requires its TRFH domain, which promotes homodimerization, DNA wrapping and recruitment of various telomere effector proteins, including Apollo, BLM and RTEL1^12,13,19–21^. At the molecular level, TRF2 binds telomeric double-stranded DNA via its Myb domain^2–4^, while its TRFH domain mediates dimerization and induces topological stress that favours invasion of the 3′-overhang into duplex telomeric DNA, generating T-loops^13,18,22,23^.

The N-terminal basic domain of TRF2 binds branched DNA junctions at the point of strand invasion of the T-loop, blocking branch migration and preventing pathological processing by resolvases, while also antagonizing PARP1 recruitment^24–27^. Additionally, TRF2 recruits cofactors such as the exonuclease Apollo to ensure proper 3′ overhang processing, which is a prerequisite for T-loop invasion, and coordinates the RTEL1 helicase to unwind T-loops during S phase to avoid replication fork collisions^19,28^. Within shelterin, TRF2 also engages RAP1 and utilizes its iDDR domain to repress NHEJ, providing a T-loop–independent layer of end-protection^29,30^. Hence, TRF2 acts both to stabilize T-loops and to directly inhibit DNA damage signalling, shielding chromosome termini from ATM activation and NHEJ.

In vitro studies with purified human TRF2 and “model telomere” DNA substrates observed that substrates containing a 3’ telomeric overhang form loops by EM^13^. Cross-linking provided evidence that the ssDNA overhang underwent strand invasion with TRF2 located at the single-double strand junction. If the overhang was removed, had discontinuities or was non-telomeric in sequence, then the loop did not form. This work established that TRF2 can promote invasion of a telomeric ssDNA overhang into dsTTAGGG repeats^13^. While evidence implicates TRF2 in stimulating T-loop formation in vitro and in cells, the precise biochemical mechanism by which it restructures telomeric DNA to form T-loops remains unclear.

Here, we have developed single-molecule assays to directly visualise shelterin activity on double-stranded telomeric DNA substrates. We show that shelterin promotes DNA reorganisation and loop formation *in vitro* through tethering of distal telomeric dsDNA repeats, which under our conditions is TRF2-dependent, involves multiple interfaces within TRF2 with the TRFH domain making a major but non-exclusive contribution, and occurs independent of the 3’ overhang. Our results support a revised model of T-loop formation, in which telomeric dsDNA tethering is a candidate first step preceding invasion of the 3’ overhang, which is essential for telomere end-protection and genome stability.

## Results

### Reconstituted human shelterin forms a stable complex containing both TRF1 and TRF2

To investigate the mechanism of shelterin-mediated chromosome end-protection we reconstituted the full human shelterin complex using the GoldenBac system^31^ (Figures 1A and S1A). Affinity tag purification and size exclusion chromatography from insect cell lysates enabled isolation of a stable complex (Figures 1B, 1C, S1B). Absorption measurements (A_260_/A_280_) and AFM of purified shelterin complex alone confirmed the absence of contaminating DNA, and the cleanliness of the protein preparations (Figures 1B, S1C and S1D).

Next, we performed mass-photometry, which confirmed the presence of ∼70% high MW (≥ 440 kDa) complexes consistent with stoichiometries of TRF1:TRF2:RAP1:TIN2:TPP1:POT1 from 2:2:2:1:1:1 (481 kDa) to 2:2:2:2:2:2 (646 kDa) (Figure 1D). Lower molecular weight species (< 440 kDa) make up the remaining 30% of the population. The identities of these species cannot be explicitly determined but are consistent with shelterin sub-complexes (e.g. previously reported TRF2:RAP1:TIN2:TPP1:POT1 at 2:2:1:1:1 stoichiometry, 368.9 kDa expected mass^32^) (Figure 1D). Western blot analysis confirmed POT1, RAP1, TPP1, TIN2 and TRF1 and/or TRF2 were present in the complex (Figure 1E).

To probe the interactivity of proteins within the purified shelterin complex, and further confirm the presence of both TRF1 and TRF2, we performed crosslinking mass-spectrometry in the absence of telomeric DNA (Figure S1E). Analysis of crosslinked peptides revealed extensive interactions between all six shelterin subunits, further confirming the reconstitution of a stable full human shelterin complex containing both TRF1 and TRF2 (Figure 1F).

Many expected interactions were observed, including specific interactions between TRF2, RAP1 and TIN2^4,33,34^; TRF1 and TIN2^8,34^, and TIN2, TPP1 and POT1^35,36^ (Figure 1F). Several unreported direct interactions were also observed between TRF1 and TRF2, confirming the presence of both TRF1 and TRF2 within the same complex. As expected additional direct interactions were observed with the full shelterin complex that were not detected by SAXS analysis of a subcomplex of shelterin comprising TRF2-RAP1 alone^37^.

To determine whether the purified shelterin complex specifically binds telomeric DNA we performed electrophoretic mobility shift assays (EMSAs). EMSAs revealed increased binding to dsDNA containing telomeric repeats compared to non-telomeric repeat sequences (Figures S1F and 1H). Competition assays confirmed a strong binding preference of shelterin to substrates combining four double-stranded telomeric repeats and at least two single-stranded telomeric repeats (Figures S1G and 1I). Binding and competition assays were also performed on agarose gels, which allow the bound complex to enter the gel and resolve as a discrete species rather than being retained in the well, and gave the same result (Figures 1G, S1H, and S1I). These results are consistent with previously reported binding affinities of TRF1 and TRF2 for telomeric dsDNA and of POT1 for telomeric ssDNA and reconstituted shelterin containing TRF2 only^32^. Together this demonstrates that reconstituted shelterin has specific affinity for telomeric DNA, and an increased affinity for telomeric DNA substrates containing a ssDNA telomeric overhang.

### Shelterin binding to telomeric DNA under applied force

Next, we developed a single-molecule optical-tweezers assay to directly observe shelterin binding to telomeric dsDNA repeats in real-time. Shelterin was fluorescently labelled with Alexa-647 by Sfp mediated CoA-dye conjugation at ybbr tags present on each protein within the complex (Figure S2A). Next, we used end-to-end ligation of plasmids containing telomeric repeats, to generate long dsDNA arrays containing regularly spaced blocks of telomeric repeats (TTAGGG)_n_, interspersed by non-repetitive dsDNA (Figures S2B and S2C). The length of each array is dependent on the number of ligated plasmids that form the array. Only DNA molecules in the size range of 15 - 30 µm (corresponding to ∼13 - 26 ligated plasmids) were used for single molecule experiments. These arrays, characterised by the number of repeats in each block (Tel_n_), were held between two optically trapped beads and extended to probe their mechanical properties (Figure 2A). Force-distance measurements of tethered Tel_80_ arrays followed the characteristic profile of an individual molecule of dsDNA under force^38^, as confirmed by fitting with the extensible worm-like-chain model (eWLC, Equation 1^39^, Figure S2D). 2D confocal imaging of Tel_80_ incubated with 2 nM fluorescently labelled shelterin revealed binding at regularly spaced intervals, consistent with specific binding to the blocks of telomeric dsDNA repeats but not to the intervening non-telomeric dsDNA (Figures 2B and S2E). We did not detect association with the intervening non-telomeric DNA at our time resolution (0.1 ms/pixel, 100 ms/line). One-dimensional diffusion of shelterin along non-telomeric DNA, and an extensive one-dimensional search by the TRF2 basic domain, have been directly observed using DNA curtains and DNA tightropes respectively^40,41^ and we draw no inference about target search from the present data.

**Figure 2.**
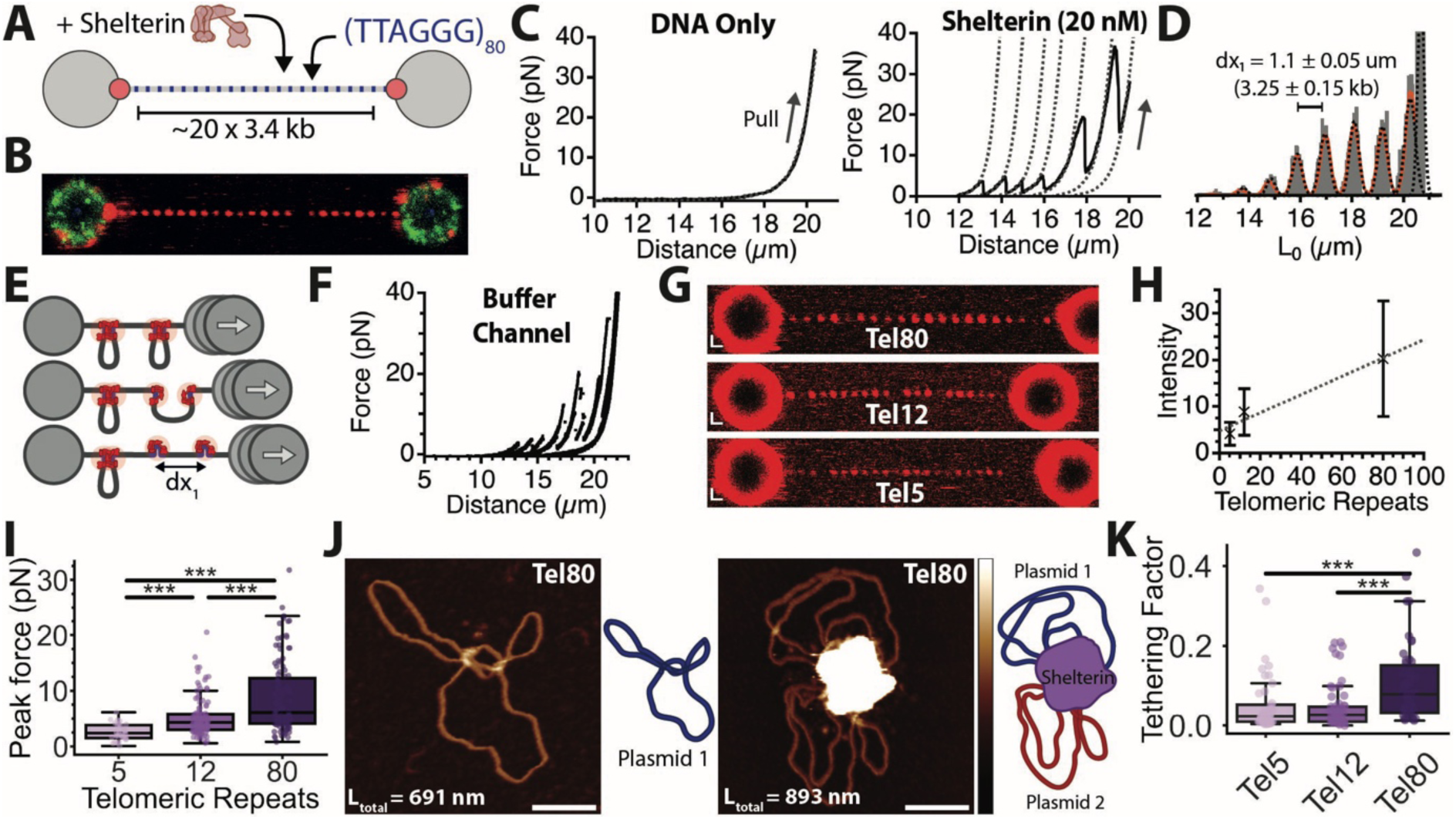
Shelterin mediates tethering between telomeric dsDNA repeats. (A) Schematic of single-molecule optical-tweezers assay. DNA arrays containing regularly spaced region of 80xTelomeric Repeats, blue, separated by non-telomeric DNA, grey, are tethered between optically trapped beads and incubated with labelled shelterin. (B) 2D confocal-microscopy image of Tel_80_ DNA arrays held at 5 pN between optically-trapped beads and incubated with 2 nM labelled shelterin (C) Force-distance curves for telomeric DNA array substrate (A) in the absence of shelterin, left, and presence of shelterin, right. Dotted lines, eWLC fit of each ‘tooth’ of saw-tooth pattern. (D) Contour length analysis for repeated cycles of DNA pulling and relaxation in the presence of shelterin. (E) Proposed model of shelterin mediated tethering of telomeric repeats. (F) Force-distance curve for DNA array incubated with shelterin before moving to buffer only channel and performing three cycles of relaxation and extension of DNA. (G) Representative 2D confocal scans of Tel_80_, Tel_12_ and Tel_5_ DNA arrays held at 5 pN and incubated with 5 nM shelterin. (H) Quantification of fluorescence intensity from shelterin bound to DNA arrays containing different length telomeric repeats. (I) Quantification of rupture forces of shelterin mediated tethering for different length telomeric repeats. Statistical significance between groups assessed with a post-hoc Dunn’s test. (J) Representative AFM image and molecule tracing of Tel_80_ plasmid imaged in the absence of shelterin, left, and presence of shelterin (5 nM), right. Scale bars = 50 nm, z scale = −2 to 5 nm. (K) Calculated normalised tethering factor of shelterin bound molecules for Tel_5_, Tel_12_ and Tel_80_ plasmids incubated with shelterin (5 nM). N values: Tel_5_ = 48, Tel_12_ = 60, Tel_80_ = 50. Statistical significance between groups assessed with a post-hoc Dunn’s test.

### Shelterin mediates tethering of distant telomeric dsDNA in a reversible manner

To determine whether shelterin binding alters telomeric DNA organisation, force-distance curves were generated for the Tel_80_ arrays in the absence or presence of shelterin (Figure 2C). In the absence of shelterin the force-distance curve followed the expected profile for dsDNA and could be fit to a single eWLC fit (L_c_ = 20.23 ± 0.01 µm, L_p_ = 50 nm, S_t_ = 1500 pN) (Figure 2C, left). Notably, on incubation with shelterin, a saw-tooth profile was observed in the force-distance measurements (Figure 2C, right). The individual sections of this saw-tooth profile could be fit with the same worm-like chain model parameters as the original DNA molecule but with a reduced DNA contour length (Figure 2C, right, dashed lines). This suggests that shelterin binding results in transient shortening of the DNA molecule, which is followed by release of a region of DNA at sufficient force. We term these release events ‘rips’, which were observed to occur at the same locations through multiple cycles of pulling and relaxation (Figure S2F, left).

To further quantify these ‘rips’ we calculated the corresponding DNA contour lengths for every data point in the force distance curves (Equation 2, L_p_ = 50 nm, S_t_ = 1500 pN; Figure S2F, right). These DNA contour length calculations revealed regularly spaced discrete peaks in observed DNA lengths which were separated by 1.1 ± 0.05 µm (Figures 2D and S2F). This corresponds to ∼3.25 ± 0.15 kb which is approximately the distance between blocks of telomeric dsDNA repeats in the arrays (Figure 2A). Each ‘rip’ in the force distance curve corresponds to the forced disruption of a single shelterin-mediated interaction between distal blocks of telomeric repeats. This data indicates that shelterin can hold together spatially distinct regions of telomeric dsDNA, which would lead to looping out of intervening DNA (Figure 2E). We term this activity ‘tethering’, defined as DNA-protein mediated DNA bridging between telomeric repeat DNA.

To probe whether shelterin-mediated tethering occurs between pre-bound shelterin complexes or requires a new binding event, telomeric arrays were incubated with shelterin with the dsDNA array extended to 10 pN, allowing binding but preventing any potential tethering activity. The dsDNA array was then moved to a buffer channel containing no shelterin complex. When the beads were brought together and the dsDNA array re-extended, tethering events were observed in the force-distance curve (Figure 2F). This demonstrates that tethering occurs between pre-bound shelterin complexes and is reversible and repeatable.

### The strength of shelterin-mediated tethering is dependent on telomere repeat length

To determine whether the strength of these tethering interactions is dependent on the number of shelterin complexes present, we generated dsDNA arrays containing different numbers of telomeric repeats (Tel_n_). On incubation with shelterin binding was observed at regularly spaced intervals for Tel_5_ and Tel_12_ arrays, corresponding to the telomeric dsDNA repeats, as observed previously for Tel_80_ (Figures 2G and S2G). Quantification of the fluorescence intensity for the different repeat lengths showed an increase in intensity with increasing number of telomeric repeats (Figures 2H and S2G). This implies that the number of shelterin molecules bound is limited by the number of telomeric dsDNA repeats.

Tethering was observed with dsDNA arrays containing all three telomere repeat lengths, Tel_5_, Tel_12_ and Tel_80_ (Figure S2H), with a significant increase in force required to disrupt tethering as the number of telomeric dsDNA repeats increased (Figures 2I and S2H). This suggests that multiple shelterin complexes bound to the telomeric dsDNA repeats contribute to the stability of tethering, which increases the force required to disrupt the interaction. This is consistent with the behaviour of previously described non-specific DNA-bridging proteins such as H-NS^42^.

### Shelterin can mediate intermolecular tethering

To determine whether telomeric dsDNA from different molecules could be tethered by shelterin, and to understand the conformational changes this drives, we employed atomic force microscopy (AFM). AFM was used to resolve the structure of a 1.92 kb pICOz plasmid containing telomeric repeats, Tel_80_, in the presence and absence of shelterin. In the absence of shelterin, high-resolution AFM images showed long supercoiled molecules formed of dsDNA (Figure 2J, left). Using an automated AFM image tracing pipeline, DNA contour length measurements of Tel_80_ were calculated as 680 ± 60 nm, consistent with the expected plasmid length of 1.92 kb (Figure 2J, left). In the presence of shelterin, we observed several plasmid DNA molecules held together by a large globular protein complex (Figure 2J, right). Because bound protein obscures a variable proportion of the underlying DNA in AFM images, the measured DNA length underestimates the amount of DNA present within a complex. Nonetheless, the total measured DNA length of shelterin-bound complexes was both greater and considerably more dispersed than that of free plasmid (753 ± 340 nm versus 680 ± 60 nm), with a subset of complexes exceeding the length expected for a single plasmid, indicating that shelterin can tether disconnected telomeric dsDNAs in trans (Figure 2J). Further AFM imaging of shelterin incubated with pICOz Tel_5_, Tel_12_ and Tel_80_ plasmids containing different length telomeric repeats regions revealed multiple plasmids tethering with all three telomere repeat lengths (Figure S2I). To better quantify tethering by AFM we developed a deep learning image segmentation pipeline to measure the DNA length and Protein volume per molecule (Figure S2J) and calculated a normalised tethering factor (DNA length x Protein volume) (Figure 2K). Bound protein ‘hides’ some of the dsDNA so we could not use DNA length alone as a measure of tethering. In agreement with the optical-tweezers experiments, the normalised tethering factor increased with an increased length of telomeric repeats (Figure 2K). This further supports the optical-tweezers data, demonstrating that shelterin-mediated tethering is dependent on telomeric dsDNA repeat length.

### TRF2 alone is sufficient for telomeric DNA tethering

To determine which subunit, or subunits, within the shelterin complex confer tethering activity, we tested TRF1 and TRF2 individually, as they possess intrinsic telomeric dsDNA binding. Tel_80_ DNA arrays incubated with TRF1 showed no tethering in force-distance curves (Figures 3A, left, and S3A), despite TRF1 binding to the telomeric repeats (Figure 3A, right). Strikingly, on incubation with TRF2 tethering was observed in the force-distance curves (Figures 3B and S3A), indicating that TRF2 alone is sufficient to drive telomeric dsDNA tethering.

**Figure 3.**
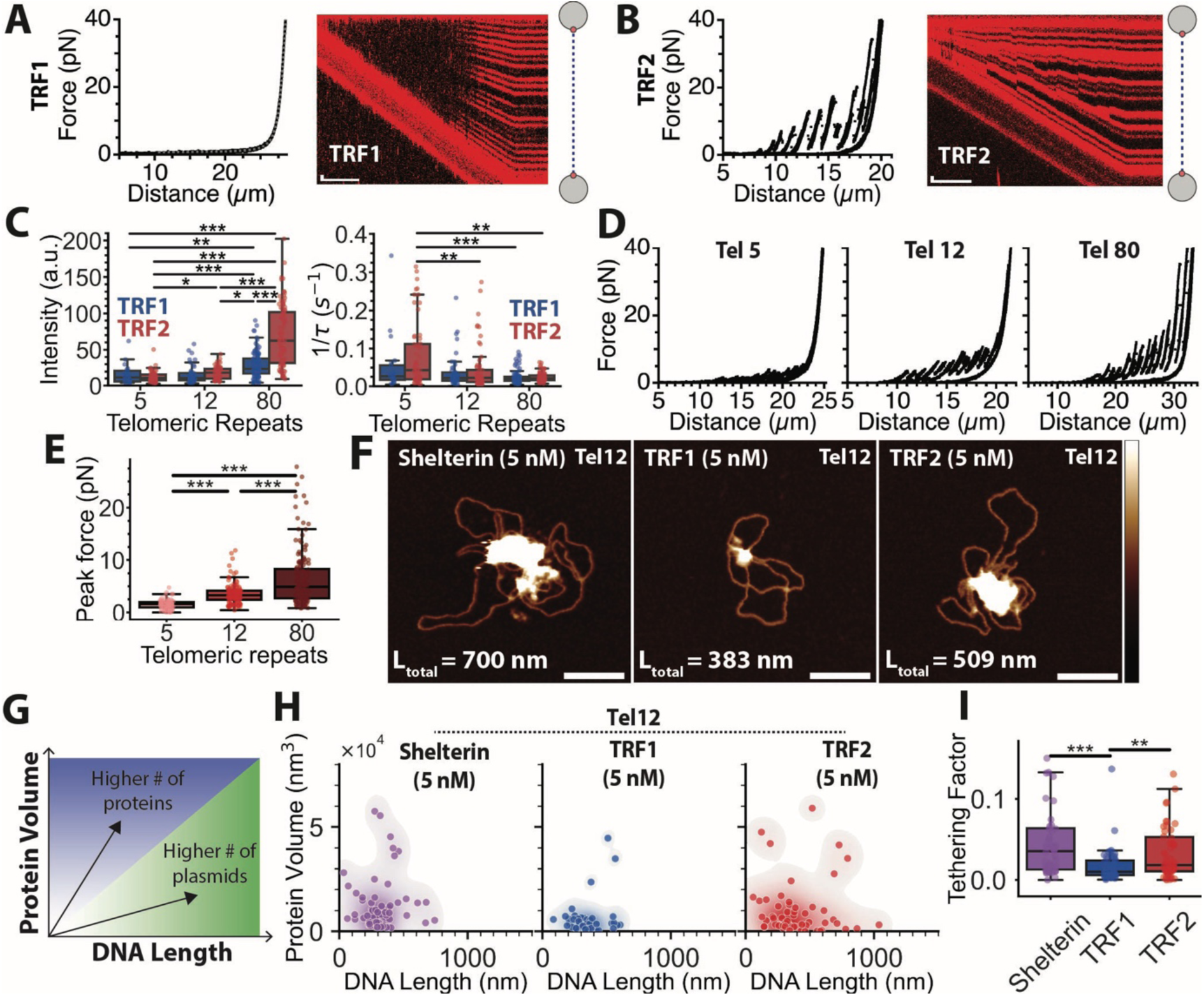
TRF2 alone is sufficient to mediate telomeric dsDNA tethering. (A) Force-distance curve, left, and corresponding kymograph, right, for TRF1 incubated with telomeric DNA array. Scale bars: x = 5 s, y = 1 µm. (B) Force-distance curve, left, and corresponding kymograph, right, for TRF2 incubated with telomeric DNA array. Scale bars: x = 5 s, y = 1 µm. (C) Quantification of initial fluorescence intensity (A.U.) of TRF1 and TRF2 bound to different length telomeric repeats, left. Quantification of dissociation rate of TRF1 and TRF2 bound to different length telomeric repeats, right. Data points are individually quantified binding sites from at least three different optically trapped DNA arrays. Statistical significance between groups assessed with a post-hoc Dunn’s test. (D) Representative force-distance plots for Tel_5_, Tel_12_ and Tel_80_ DNA array constructs incubated with TRF2. (E) Quantification of average rupture force as a function of number of telomeric repeats in the DNA array. Data points are individual rupture events from at least three different optically trapped DNA molecules. Statistical significance between groups assessed with a post-hoc Dunn’s test. (F) Representative AFM images for Tel_12_ telomeric repeat containing plasmid incubated with shelterin, TRF1 and TRF2 (5 nM). Scale bars = 50 nm, z scale = −2 to 5 nm. L_total_ = total measured DNA length (G) Schematic of protein volume vs total DNA length analysis plots (H) Quantification of protein volume vs total DNA length for Tel_12_ plasmids incubated with shelterin, TRF1 and TRF2 (5 nM). (I) Comparison of calculated normalized tethering factor from AFM measurements of Tel_12_ plasmids incubated with shelterin, TRF1 and TRF2 (5 nM). N values: shelterin = 60, TRF1 = 61, TRF2 = 62. Statistical significance between groups assessed with a post-hoc Dunn’s test.

To determine whether TRF2-mediated tethering shows the same telomere repeat length dependence as shelterin, Tel_5_, Tel_12_ and Tel_80_ dsDNA arrays were incubated with TRF1 and TRF2. We first quantified the initial fluorescent intensity and dissociation rate of TRF1 and TRF2 bound to Tel_5_, Tel_12_ and Tel_80_ dsDNA arrays. We observed a higher initial fluorescence intensity with increasing length of telomeric repeats, and higher intensity of TRF2 observed with Tel_80_ arrays than TRF1 (Figures S3B, S3C and 3C, left). We next quantified the dissociation rates of TRF1 and TRF2 (Figures S3B, S3C and S3D). An overall decrease in dissociation rate was observed for both TRF1 and TRF2 as the length of telomeric repeats increased, with no significant differences in rate observed between TRF1 and TRF2 (Figure 3C, right).

Tethering was observed with TRF2 for all three telomere repeat lengths (Figure 3D). Analysis of the rupture forces revealed a significant increase in force required to disrupt the TRF2-mediated tethering as the number of telomeric repeats increases (Figure 3E). Together, this suggests that, like the full shelterin complex, multiple DNA bound TRF2 molecules contribute additively to the tethering of distal telomeric dsDNA regions.

### Shelterin and TRF2, but not TRF1, form complexes that tether dsDNA molecules

To understand how the tethering activity of the full shelterin complex compares to that of TRF1 or TRF2, we used AFM to probe differences in the bound protein and DNA. We observed protein binding to Tel_12_ plasmids for TRF1, TRF2 and shelterin as expected (Figure 3F). Using our automated image segmentation pipeline, we quantified the protein volume and length of DNA within bound complexes for each species (Figures 3G and 3H) and calculated the normalised tethering factor for each molecule (Figure 3I). Consistent with the optical-tweezers experiments, TRF2 showed levels of plasmid tethering activity comparable to shelterin and significantly higher than TRF1 (Figure 3I). We performed additional experiments on Tel_5_ at 5 nM protein concentrations (Figures S3E and S3F) and on Tel_12_ with two higher protein concentrations: 10 nM (Figures S3G and S3H), and 20 nM (Figures S3I and S3J). We consistently observed a lack of tethering by TRF1 compared to shelterin and TRF2 (Figures S3E – S3J). Based on our comparison of Tel_5_, Tel_12_ and Tel_80_ by both optical-tweezers and AFM we determined the Tel_12_ repeat length to be the optimal substrate for quantitative analysis, enabling the most accurate quantification and the best dynamic range to detect changes in activity.

Together this data supports the optical-tweezers data, demonstrating that shelterin and TRF2 have comparably high levels of telomeric dsDNA tethering activity compared to TRF1. In contrast, TRF1 does not show tethering activity by optical-tweezers and shows significantly reduced tethering activity by AFM.

### Tethering is not uniquely dependent on the TRF2 TRFH domain or TRF2 dimerization

As the TRFH domain of TRF2 has been previously demonstrated to mediate TRF2 dimerization and is essential for T-loop formation *in vivo*^43^, we hypothesised that the TRFH domain may be required for tethering. To test this possibility, we purified and fluorescently labelled a TRF2 TRFH domain deletion mutant, TRF2ΔTRFH (Figure S4A). Next, we performed mass photometry to determine the dimerization state of the deletion mutant (Figure 4A). For TRF1 and TRF2, a clear peak at the expected molecular weight of a dimer was observed (TRF1: 110 ± 5 kDa; expected = 110 kDa, TRF2: 117 ± 7 kDa; expected = 121 kDa). However, for TRF2ΔTRFH, a single peak corresponding to the expected molecular weight of a monomer was observed (TRF2ΔTRFH: 37 ± 4 kDa; expected = 37 kDa) (Figure 4A). This confirms that deletion of the TRFH domain from TRF2 abolishes dimerization in solution.

**Figure 4.**
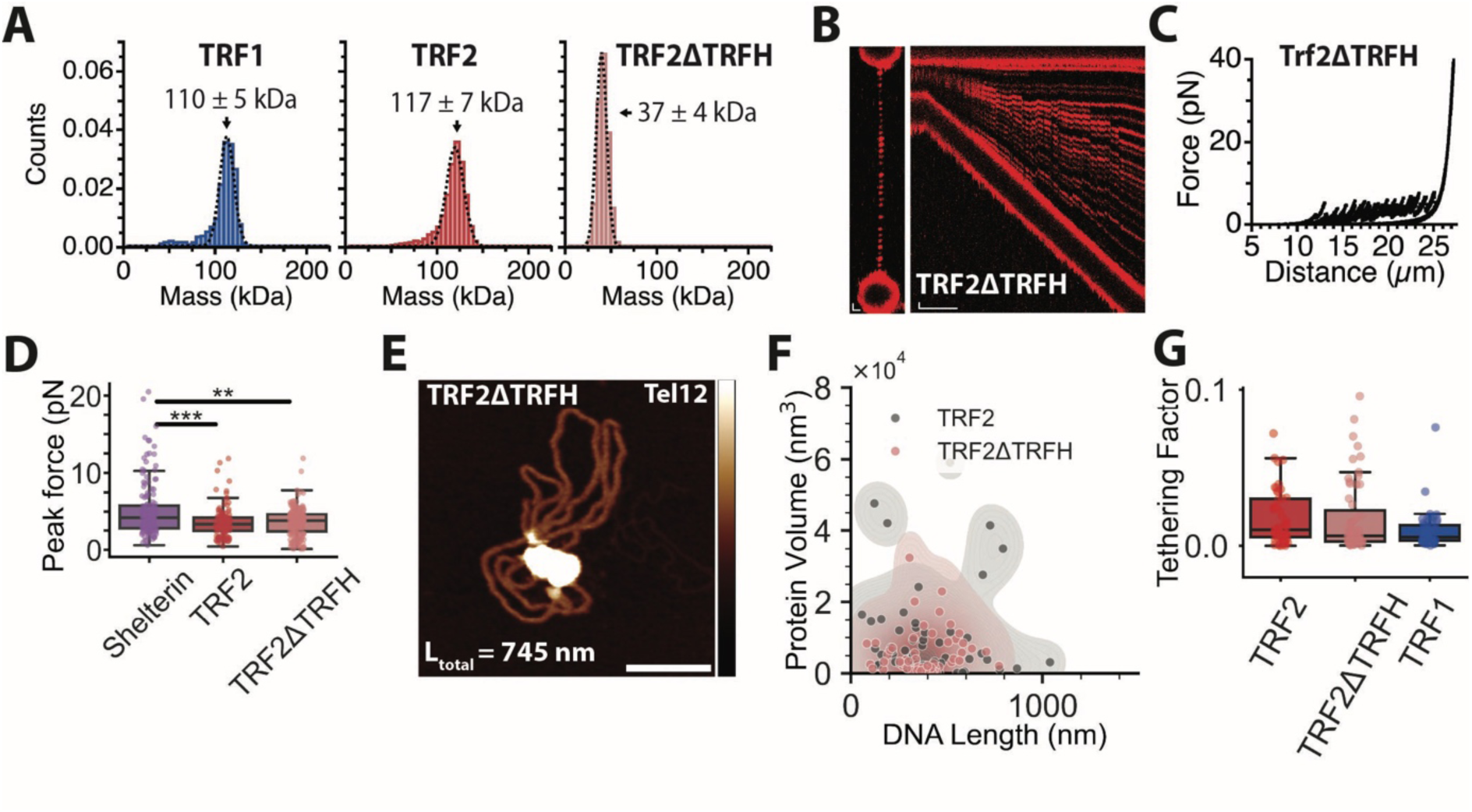
Telomeric dsDNA tethering does not strictly require the TRF2 TRFH domain or TRF2 dimerization. (A) Mass photometry of TRF1, TRF2 and TRF2ΔTRFH showing stable dimer of TRF1 and TRF2 and monomer of TRF2ΔTRFH. (B) Representative 2D confocal scan of Tel_12_ DNA array construct incubated with TRF2ΔTRFH, left, and kymograph of force-distance measurement, right. 2D scan scale bars = 1 µm; kymograph scale bars: x = 5 s, y = 1 µm. (C) Representative force-distance measurement of Tel_12_ DNA array constructs incubated with TRF2ΔTRFH. (D) Comparison of rupture forces for Tel_12_ DNA array construct incubated with shelterin, TRF2 and TRF2ΔTRFH. Data points are individual rupture events from across at least three different optically trapped DNA arrays. Statistical significance between groups assessed with a post-hoc Dunn’s test. (E) Representative AFM image of plasmid containing telomeric repeats incubated with TRF2ΔTRFH. Scale bars = 50 nm, z scale = −2 to 5 nm. (F) Quantification of protein volume vs total DNA length observed by AFM for Tel_12_ plasmids incubated with TRF2, grey, and TRF2ΔTRFH, red. (G) Comparison of calculated normalized tethering factor from AFM of Tel_12_ plasmid incubated with TRF2 and TRF2ΔTRFH. N values: TRF2 = 62, TRF2ΔTRFH = 63, TRF1 = 61. Statistical significance between groups assessed with a post-hoc Dunn’s test.

To determine whether loss of TRF2 dimerization affects binding and tethering activity, we incubated TRF2ΔTRFH with Tel_12_ array telomeric repeat-containing dsDNA arrays (Figure 4B). Specific binding to telomeric repeats was observed across the DNA (Figure 4B, left). On DNA relaxation and extension, ‘rips’ were observed in the force-distance curve (Figure 4C). Analysis of the rupture forces showed that TRF2ΔTRFH retains telomeric dsDNA tethering activity comparable to TRF2 (Figure 4D).

Next, we performed AFM experiments, which revealed binding and telomeric dsDNA tethering by TRF2ΔTRFH when incubated with Tel_12_ (Figures 4E and 4F). Quantification and calculation of the tethering factor of TRF2ΔTRFH revealed an activity between that of TRF1 and TRF2 (Figures 4F and 4G). This suggests that while the TRFH domain may contribute to the tethering activity of TRF2, its deletion does not eliminate tethering activity, raising the possibility that more than one region of TRF2 contributes to tethering activity.

To understand whether tethering activity could be due to increased affinity for telomeric repeats we used fluorescence polarisation to measure the specific binding affinity of TRF1, TRF2 and TRF2ΔTRFH for DNA containing a four telomeric repeat binding site (Figure S4B, Table 1). TRF1 displayed the highest specific affinity for binding telomeric repeats, K_d_ = 0.9 ± 0.9 nM, at the tight-binding limit of the assay, and substantially higher than that of TRF2, K_d_ = 8.3 ± 2.0 nM. These K_d_ values are consistent with previously reported literature, where similar salt and pH conditions where used (shelterin core with dsDNA/ssDNA telomeric overhang, K_d_ = 1.1 ± 0.2 nM ^32^; TRF1 myb domain only, telomeric dsDNA, K_d_ = 3.2 ± 0.5 nM^44^; TRF2, K_d_ 10.3 ± 2.9 nM^45^). Notably, the TRF2ΔTRFH protein retained a high specific affinity for telomeric repeats (K_d_ = 16.9 ± 1.1 nM), albeit lower than TRF2. The observed 2-fold lower specific binding affinity could be due to loss of its ability to dimerize. These measurements demonstrate that the tethering activity observed for TRF2 and TRF2ΔTRFH, but not TRF1, is not driven by higher affinities for binding telomeric repeats.

**Table. 1.** Binding affinities (K_d_) and Hill coefficients (n_H_) for specific binding to telomeric DNA measured by fluorescence polarisation.

| Protein | K<sub>d</sub> (nM) | n<sub>H</sub> |
| --- | --- | --- |
| TRF1 | 0.9 ± 0.9 | 3 ± 8 |
| TRF2 | 8.3 ± 2 | 3 ± 4 |
| TRF2ΔTRFH | 16.9 ± 1.1 | 6 ± 6 |
| TRF2cA<sub>TRF1</sub> | 1.1 ± 0.2 | 3 ± 2 |
| TRF2cTRFH<sub>TRF1</sub> | 2.5 ± 0.6 | 3 ± 2 |
| TRF2cHinge<sub>TRF1</sub> | 1.4 ± 0.4 | 2 ± 1 |
| TRF2cMyb<sub>TRF1</sub> | 7.6 ± 1.4 | 7 ± 8 |
| TRF1cTRFH<sub>TRF2</sub> | 4.7 ± 1.3 | 2 ± 1 |

### The TRF2 TRFH domain contributes to telomeric DNA tethering but is not the sole driver

To further investigate the basis of TRF2-mediated tethering, we generated chimeric proteins of TRF2 containing domains swapped with the corresponding TRF1 domains (Figure S4C). To confirm that all domain swap mutants retained activity for binding telomeric repeats fluorescence polarisation was used to measure binding affinities (Figure S4D). All domain swap mutants retained high specific affinity for telomeric repeats. Hill coefficients (n_H_) are reported in Table 1 as fitting parameters only; the associated uncertainties are too large to support an interpretation of binding cooperativity. Interestingly, TRF2cAcidic_TRF1_, TRF2cTRFH_TRF1_ and TRF2cHinge_TRF1_ domain swap mutants displayed tighter DNA binding affinities than wtTRF2, with affinities comparable to that of TRF1 (Table 1). In contrast, the binding affinity of TRF2cMyb_TRF1_ was comparable to that of TRF2 (Table 1). This suggests the increased DNA binding affinity of TRF1 is due to domains other than the Myb domain.

We next tested these domain swap mutants for tethering activity. We reasoned that when a domain from TRF2, which confers tethering, was swapped with the corresponding domain from TRF1, tethering activity would be lost. Interestingly, on incubation with Tel_80_ dsDNA arrays, all domain swap mutants of TRF2 with domains from TRF1 retained tethering activity (Figure S4E). Notably, deletion of the TRF2 Basic domain (TRF2ΔBasic) also retained tethering (Figure S4F, left), whereas TRF2ΔMyb lacked tethering as it is unable to bind telomeric dsDNA (Figure S4F, right). This data raised the possibility that tethering is a multi-valent interaction involving several domains across TRF2. This is consistent with cross-linking mass-spectrometry data for recombinant shelterin, which revealed multiple internal inter-domain cross-links for TRF2 (Figure S4G).

Next, we reasoned that if the TRFH domain of TRF2 contributes to tethering, as observed in the AFM measurements of TRF2ΔTRFH (Figure 4G), swapping the TRFH domain from TRF2 into TRF1 should confer tethering activity to TRF1. To test this possibility, we generated a reciprocal domain swap mutant, TRF1cTRFH_TRF2_ (Figure 5A). TRF1cTRFH_TRF2_ displays an intermediate DNA binding affinity between that of TRF1 and TRF2, as measured by fluorescence polarisation (Table 1, Figure S4H). On incubation of TRF1cTRFH_TRF2_ with Tel_12_ telomeric repeat-containing DNA, characteristic ‘rips’ were observed in the force-distance curves (Figure 5B). This demonstrates that the TRFH_TRF2_ domain is sufficient to confer tethering activity to TRF1. We also observed tethering activity for TRF2cTRFH_TRF1_ (Figure 5C), consistent with our observation that TRF2ΔTRFH retains tethering activity (Figure 4). Rupture forces required to disrupt tethering by TRF1cTRFH_TRF2_ and TRF2cTRFH_TRF1_ were comparable to wild type TRF2 (Figure 5D).

**Figure 5.**
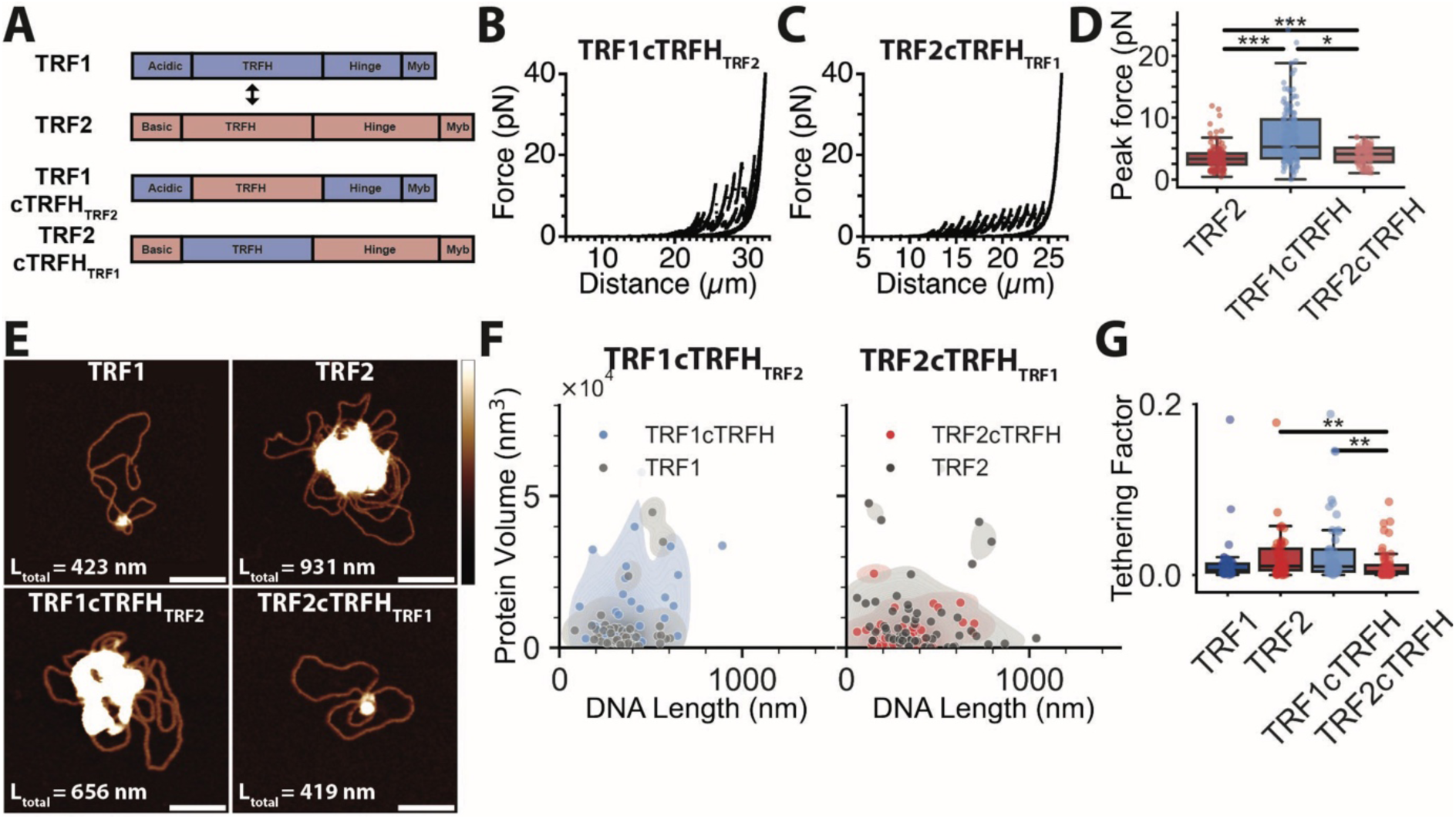
The TRF2 TRFH domain contributes to telomeric DNA tethering but is not solely responsible for it. (A) Schematic of domain swap variants between TRF1 and TRF2. (B) Representative force-distance measurement of Tel_12_ DNA array constructs incubated with TRF1cTRFH domain swap mutant. (C) Representative force-distance measurement of Tel_12_ DNA array constructs incubated with TRF2cTRFH domain swap mutant. (D) Comparison of rupture forces for Tel_12_ DNA array construct incubated with TRF2, TRF1cTRFH_TRF2_ and TRF2cTRFH_TRF1_. Data points are individual rupture events from across at least three different optically trapped DNA arrays. Statistical significance between groups assessed with a post-hoc Dunn’s test. (E) Representative AFM images of Tel_12_ plasmid incubated with TRF1, TRF2, TRF1cTRFH_TRF2_ and TRF2cTRFH_TRF1_. Scale bars = 50 nm, z scale = −2 to 5 nm. (F) Quantification of protein volume vs total DNA length observed by AFM for Tel_12_ plasmids incubated TRF1 (grey) and TRF1cTRFH_TRF2_ (blue), left, and TRF2 (grey) and TRF2cTRFH_TRF1_ (red), right. (G) Comparison of calculated normalized tethering factor from AFM of Tel_12_ plasmid incubated with TRF1, TRF2, TRF1cTRFH_TRF2_ and TRF2cTRFH_TRF1_. N values: TRF1 = 61, TRF2 = 62, TRF1cTRFH_TRF2_ = 62, TRF2cTRFH_TRF1_ = 62. Statistical significance between groups assessed with a post-hoc Dunn’s test.

We further investigated the tethering activities of the TRF1 and TRF2 TRFH domain swap mutants by AFM (Figure 5E). For TRF1cTRFH_TRF2_ we observed large volumes of protein associated with ∼2 plasmid DNA molecules, similar to wild type TRF2. For TRF2cTRFH_TRF1_ we observed reduced protein binding, usually to a single plasmid DNA molecule (Figure 5E). Quantitative analysis and calculation of normalised tethering factor confirmed that TRF2cTRFH_TRF1_ exhibits significantly lower tethering activity than TRF2 (Figures 5F and 5G). In contrast, TRF1cTRFH_TRF2_ shows significantly increased tethering activity compared with TRF1 and TRF2cTRFH_TRF1_, confirming that the TRFH domain of TRF2 can promote tethering of telomeric dsDNA molecules. Together these results show that the TRFH domain of TRF2 is sufficient for telomeric dsDNA tethering but is not exclusively required.

### The TRF2 TRFH domain mediates capture of telomeric DNA in trans

Next, we used optical tweezers to test whether TRF1 and TRF2 could capture non-telomeric or telomeric dsDNA *in trans* (Figure 6A). We used fluorescently labelled DNA constructs lacking a ssDNA overhang to allow us to specifically probe dsDNA capture activity rather than invasion of the 3’ telomeric overhang. Tel_12_ DNA arrays were incubated with saturating concentrations of TRF1 (10 nM) and labelled dsDNA oligos containing either four scrambled telomeric repeat sequences (4xNoTel) or four intact telomeric repeats (4xTel) (Figure 6B). Minimal capture of the labelled non-telomeric or telomeric dsDNA was observed (Figure 6B). Next, Tel_12_ DNA arrays were incubated with saturating concentrations of TRF2 (10 nM). On addition of the labelled non-telomeric dsDNA a low level of capture was observed (Figure 6C, Top). However, incubation with the telomeric substrate, revealed high levels of dsDNA capture (Figure 6C, Bottom). Quantitative fluorescent analysis of the kymograph line intensities confirmed low levels of TRF2 capture of non-telomeric dsDNA, and dramatically higher levels of TRF2 mediated telomeric DNA capture compared to TRF1 (Figure 6D). This data again demonstrates a specific activity of TRF2 in tethering telomeric dsDNA.

**Figure 6.**
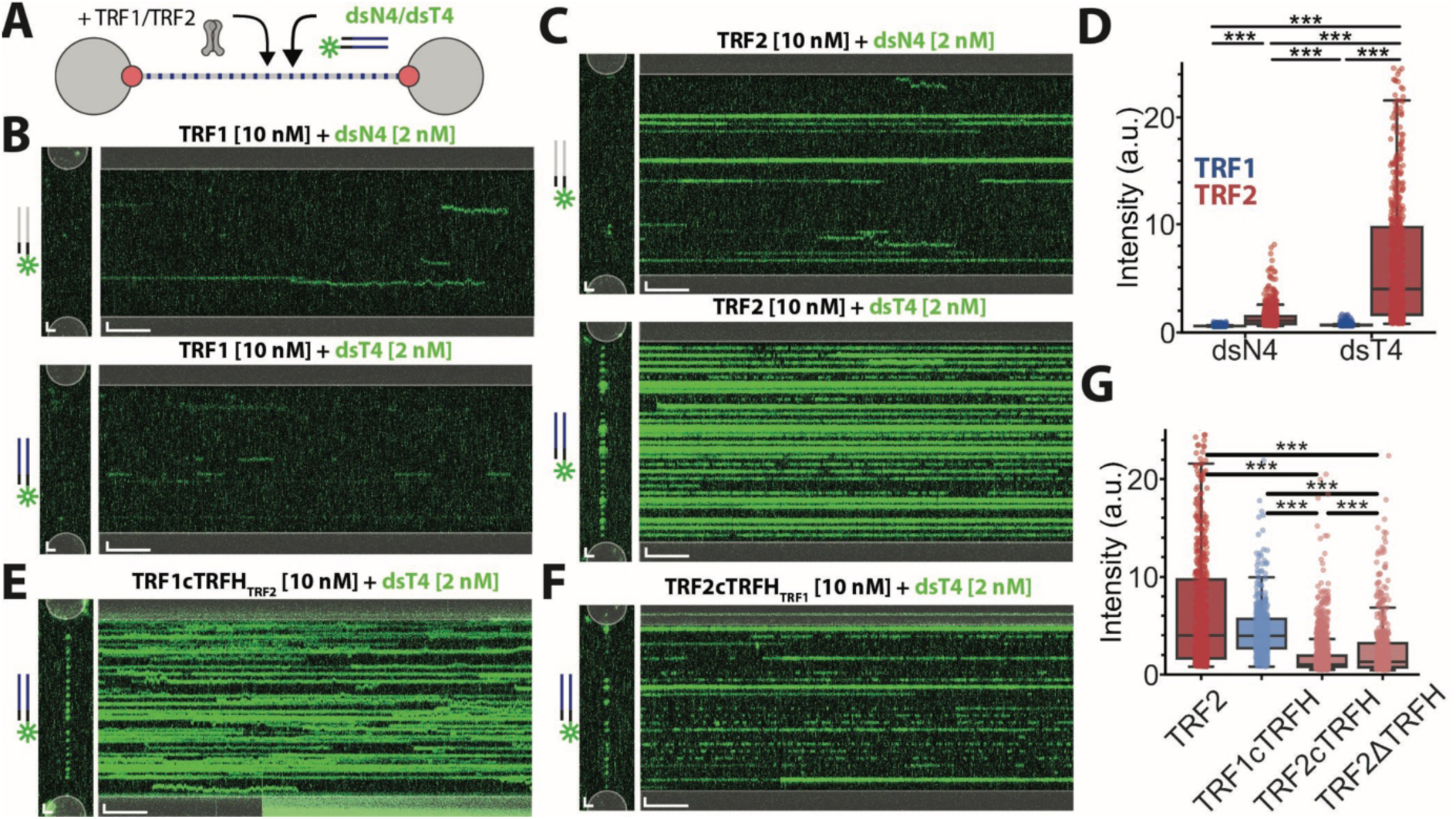
TRF2 TRFH domain mediates capture of telomeric DNA in trans. (A) Schematic of trans capture assay. Tel_12_ DNA arrays are incubated with unlabelled telomeric proteins and fluorescently labelled dsDNA containing either four telomeric repeats (4xTel) or four scrambled telomeric sequences (4xNoTel). (B) 2D-scans and corresponding kymographs for Tel_12_ DNA arrays incubated with TRF1 (10 nM) and labelled 4xNoTel dsDNA (2 nM), top, or 4xTel dsDNA (2 nM), bottom. (C) 2D-scans and corresponding kymographs for Tel_12_ DNA arrays incubated with TRF2 (10 nM) (D) Quantification of line intensity across kymographs recorded for trans capture assays with TRF1 and TRF2 in the presence of labelled dsN4 (4 x scrambled telomeric repeats) or dsT4 (4 x telomeric repeats). Data points are individual line intensities spanning 10 μm of the Tel_12_ DNA array from across at least three different optically trapped DNA arrays. Statistical significance between groups assessed with a post-hoc Dunn’s test. (E) 2D-scans and corresponding kymograph for incubation of Tel_12_ DNA arrays with TRF1cTRFH_TRF2_ domain swap mutant and labelled 4xTel dsDNA. (F) 2D-scans and corresponding kymograph for incubation of Tel_12_ DNA arrays with TRF2cTRFH_TRF1_ domain swap mutant and labelled 4xTel dsDNA. (G) Quantification of line intensity across kymographs recorded for trans capture assay of telomeric dsDNA with TRF2, TRF1cTRFH_TRF2_ and TRF2cTRFH_TRF1_ and TRF2ΔTRFH. Data points are individual line intensities spanning 10 μm of the Tel_12_ DNA array from across at least three different optically trapped DNA arrays. Statistical significance between groups assessed with a post-hoc Dunn’s test. For panels B, C, E and F 2D-scan x and y scale bars = 1 μm. Kymograph y scale bars = 1 μm, x scale bars = 10 s.

Based on our previous optical-tweezers and AFM data we hypothesised that the TRFH domain should also be involved in this telomeric DNA capture activity in trans. Indeed, domain swap of the TRF1 TRFH domain with the TRF2 TRFH domain (TRF1cTRFH_TRF2_) resulted in a high level of telomeric dsDNA capture activity, comparable to the activity of wild-type TRF2 (Figures 6E, 6G and S6A). In contrast, the TRF2cTRFH_TRF1_ domain swap resulted in a significant reduction in DNA capture compared to TRF2 (Figures 6F, 6G and S6B). To further test this we performed telomeric DNA *in trans* capture experiments with TRF2ΔTRFH (Figure S6C). TRF2ΔTRFH showed a significant reduction in tethering activity compared to TRF2, with activity comparable to TRF2cTRFH_TRF1_ (Figure 6G). Together this data further supports the role of the TRF2 TRFH domain in TRF2-specific dsDNA tethering activity.

## Discussion

Telomeres are protected by the shelterin complex, which prevents chromosome ends from eliciting a DNA damage response and undergoing promiscuous end-joining. A central feature of end-protection is the formation of T-loops, in which the single-stranded telomeric overhang invades upstream duplex telomeric repeats to mask the chromosome end. While TRF2 has long been established as the sole shelterin subunit essential for T-loop formation in cells, the underlying biochemical mechanism has remained unclear. Prior work has suggested that both TRF1 and TRF2 are able to locally re-organise the DNA^45,46^. The work presented here identifies a dsDNA tethering activity intrinsic to TRF2 that brings distal telomeric repeats into close proximity prior to strand invasion of the 3’ overhang. Those earlier single-molecule studies examined compaction and local distortion within a single continuous telomeric tract, whereas the assays reported here probe bridging between telomeric tracts separated by several kilobases of non-telomeric DNA. The long DNA arrays and force regime used in our optical-tweezers experiments do not provide the force or length resolution required to detect local compaction of the kind previously reported for TRF1, so the absence of TRF1-mediated tethering in our assays is not in conflict with those observations; the two activities are operationally distinct.

By combining single-molecule optical tweezers and AFM assays, and analysing telomeric dsDNA substrates lacking the ssDNA overhang, we have discovered that the shelterin complex and TRF2 alone can tether spatially separated telomeric dsDNA repeats, with tethering strength increasing in proportion to telomeric repeat length. Our data indicate that more than one region of TRF2 contributes to this activity, although the physical basis of the bridge remains to be determined. Furthermore, TRF2 alone can specifically capture telomeric dsDNA in *trans*. Importantly, this activity is reversible and repeatable, consistent with the dynamic nature of T-loop formation and the need for transient disassembly during DNA replication.

Under the conditions used here, force-resistant tethering is TRF2-dependent: TRF1 does not support it, whereas TRF2 does. This does not imply that TRF1 is incapable of bridging DNA. TRF1-mediated bridging of telomeric DNA *in trans* has been reported, as has TRF2-TIN2-mediated bridging ^47,48^ but the TRF1 bridges described in that work are transient, with lifetimes of approximately two seconds on short telomeric substrates unless stabilised by TIN2. Our assay registers only bridges that persist for seconds to minutes and that bear loads of tens of piconewtons, so a transient, non-force-bearing bridge would not be detected. The distinction we observe between TRF1 and TRF2 is therefore one of bridge lifetime and mechanical strength rather than of the existence of bridging as such. We note also that those studies did not compare the TRF1 and TRF2 directly: Pan et al. examined TRF1 and TIN2 without TRF2, and the corresponding TRF2 study from the same laboratory used different substrates to address a different question, with the two proteins appearing together only in a competition assay scored on T-loop formation frequency. Neither applies a defined mechanical load. Here the two proteins are compared in parallel, using the same preparations and substrates, in three independent geometries: under a force ramp, where TRF1 supports no tethering despite binding the telomeric repeats; at constant applied load in the *in trans* capture assay, where TRF2 captures telomeric dsDNA at significantly higher levels than TRF1; and in the absence of applied force by AFM, where TRF1-mediated tethering is significantly reduced relative to TRF2 and shelterin. That the AFM activity is reduced rather than abolished is consistent with the transient TRF1 bridges reported previously, and we do not claim that TRF1 is incapable of bridging telomeric DNA; the difference between TRF1 and TRF2 is largest where a load is applied. This offers a biochemical basis for the long-standing question of why TRF2 is specifically required for T-loop formation, despite the two proteins having evolved from a common ancestor and sharing DNA-binding domains and a common domain structure. Notably, TRF1 binds telomeric repeats with a higher specific affinity than TRF2 (**Table 1**), so the distinction between the two paralogues reflects a distinct biochemical activity rather than differential occupancy of the substrate. The reconstituted complex tethers with an efficiency comparable to that of TRF2 alone, indicating that TIN2, TPP1, POT1 and RAP1 neither confer nor occlude this activity, and that TRF1 within the same complex does not interfere with TRF2-mediated bridging. We note that the reconstituted complex contains both TRF1 and TIN2, so a contribution from TRF1-TIN2-mediated bridging to the activity of the full complex cannot be excluded.

Domain swap and deletion analysis further refine our understanding of the tethering mechanism. The TRFH domain of TRF2 is sufficient to confer telomeric dsDNA tethering when introduced into TRF1, yet its removal does not abolish TRF2-mediated tethering, demonstrating that tethering is not mediated by a single domain. These experiments identify regions that influence tethering but do not define the physical bridging interface: the residual activity of TRF2ΔTRFH could arise from the basic region, the hinge or the Myb domain, or from DNA-induced association of multiple TRF2 molecules, and we exclude none of these possibilities. Goldfarb *et al*.^49^, have recently reported that the hinge domain of TRF2 contributes to tetramerization and T-loop formation *in vivo*, which may provide an additional interface that could contribute to TRF2-dependent tethering. Notably, replacing the TRF2 hinge with the corresponding region of TRF1 did not abolish tethering in our assays (Figure S4E), indicating that hinge-mediated higher-order assembly is not strictly required for telomeric dsDNA tethering in vitro and that tethering and tetramerization are separable activities. Dimerization has previously been reported to be required for stable telomeric DNA binding by TRF-family proteins, yet monomeric TRF2ΔTRFH retains high specific affinity (Kd = 16.9 ± 1.1 nM, approximately two-fold weaker than TRF2) and substantial tethering activity. On an array of tandem repeats, avidity arising from multiple independently engaged Myb domains would be expected to compensate in part for the loss of a constitutive dimer interface, which may explain why TRF2 tolerates loss of the TRFH domain better than has been reported for TRF1. These insights reconcile previous *in vivo* findings implicating TRFH-dependent dimerization in T-loop formation with the observation that additional TRF2 interfaces contribute to telomeric DNA remodelling^48–52^.

Based on the data presented here, we propose telomeric dsDNA tethering by TRF2 as a candidate first step in T-loop formation, and present a model involving at least three distinct steps, only the first of which is demonstrated in this work (Figure 7). 1) telomeric dsDNA repeat tethering by TRF2 brings the chromosome end in close juxtaposition with internal telomeric repeats (This work); 2) a topological transition facilitates invasion of the 3’ ssDNA overhang into telomeric dsDNA repeats (inferred from previous work; not demonstrated here), and 3) stabilisation of the T-loop junction by the TRF2 basic domain preventing branch migration and conversion to resolvase-sensitive 4-way junctions (inferred from previous work; not demonstrated here) (Figure 7).

**Figure 7.**
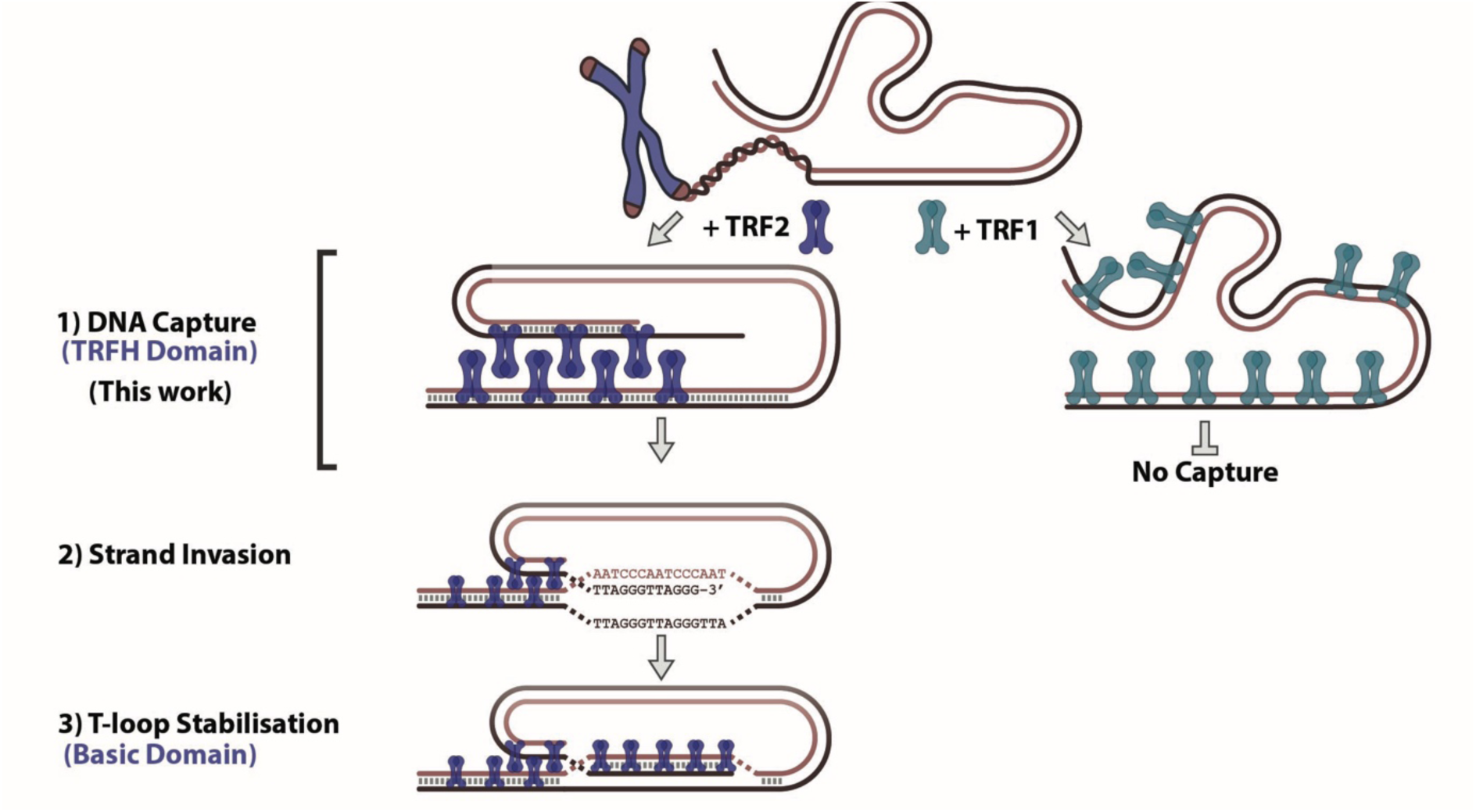
Proposed model for T-loop formation through TRF2 mediated telomeric DNA capture. We propose that TRF2 tethering activity identified in this work forms the first step in the proposed mechanism of T-loop formation. 1) DNA Capture, TRF2 tethering activity, involving the TRFH domain, holds telomere end close to internal telomeric DNA repeats (this work); 2) Strand Invasion, 3’ end of the telomeric end invades into the dsDNA (inferred from previous work, specific mechanism is currently unknown); 3) T-loop Stabilisation, TRF2 binding and specific activity of the basic domain stabilises the T-loop junction (inferred from previous work, specific mechanism is currently unknown).

Two limitations of this work should be stated explicitly. First, the substrates used here are circular and contain neither a DNA end nor a 3’ single-stranded overhang, and we measure no strand invasion, D-loop or T-loop. What we demonstrate is bridging between distal telomeric duplex repeats and the consequent looping of the intervening DNA; the relationship of this activity to T-loop formation is a model and testing it will require an end-containing substrate carrying a defined 3’ overhang together with a direct readout of strand invasion. Second, compaction within a single telomeric tract and bridging between distal tracts are distinct activities with distinct force-distance signatures, and our assays are configured to detect the latter. Compaction produces a progressive shortening of the tether, whereas the events reported here appear as an abrupt increase in extension at a defined force that restores the contour length expected for the fully extended array and that recurs at the same positions on successive pulling cycles. The absence of TRF1-mediated tethering in our experiments therefore carries no implication for TRF1-mediated compaction, which has been reported elsewhere.

By starting with dsDNA repeats alone we uncovered a tethering activity that does not require the ssDNA overhang and that may have been missed using a substrate containing both double- and single-stranded repeats; follow-up studies with such substrates will allow the transition from tethering to strand invasion to be investigated directly. Our experiments were also performed in the absence of nucleosomes, and tethering is likely to be influenced by telomeric chromatin, which differentially impacts TRF1 and TRF2 binding and displays unique structural properties^50–54^. Incorporating nucleosomes into the reconstituted system would be a powerful way to investigate the interplay between TRF2-mediated tethering, chromatin structure and T-loop formation. Finally, the domain swaps used here alter more than the tethering interface, since the TRFH domain is also the docking site for partner proteins^32^, establishing the contribution of tethering to T-loop formation in cells will require separation-of-function alleles that abolish tethering while preserving dimerization, telomeric DNA binding and partner recruitment.

Overall, these results provide direct biochemical evidence that TRF2 bridges distal telomeric duplex repeats, identifying force-resistant dsDNA tethering as an activity that distinguishes TRF2 from TRF1 under these conditions. By characterising the mechanical properties of this bridge and identifying it as a candidate first step in the remodelling of telomeric DNA into protective T-loops, this work advances our understanding of the molecular basis of chromosome end protection, which plays a fundamental role in maintaining genome stability and in averting ageing and telomere-associated disease.

## Methods

### Equations

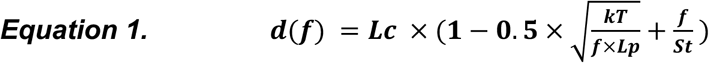

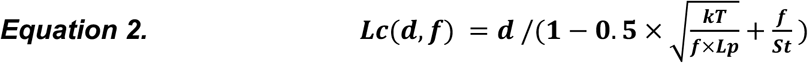

### Protein expression

The human shelterin complex and individual TRF1 and TRF2 proteins were expressed in insect cells from pFastBac1 vectors as previously described^53,54^. For TRF1, TRF2 and Rap1, constructs contain N-terminal histidine (6XHis) tags followed by ybbr-tag (DSLEFIASKLA) for fluorescent labelling using Sfp phosphopantetheinyl transferase^55,56^. For Tpp1, Tin2 and Pot1, constructs contain N-terminal 6XHis-MBP tags with PreScission sites for tag cleavage. Tpp1 and Pot1 contain N-terminal ybbr-tags and Tin2 contains a C-terminal ybbr-tag. TRF1 and TRF2 mutants (Table S2) were generated by site-directed mutagenesis or Takara In-Fusion cloning of the original vectors and verified by whole plasmid sequencing.

The vectors were used to generate baculovirus as per the Bac-to-Bac Baculovirus Expression System. Small scale expression tests were performed to determine the optimum viral titre. For shelterin expression, baculovirus for all six proteins was used to infect 1.6 liters of Sf9 insect cells at ∼2 x 10^6^ cells ml^-1^ and incubated for 72 hrs (27 °C, 120 rpm). For TRF1, TRF2 and respective mutants, 0.8 liters of Sf9 cells were infected at ∼2 x 10^6^ cells ml^-1^ and incubated for 72 hrs (27 °C, 120 rpm). Cells were collected and washed with cold PBS, snap-frozen in liquid nitrogen and stored at −80 for purification.

### Protein Purification

Full human shelterin complex was purified from insect Sf9 cells. Cells were seeded at 1×10^6 cells/ml in 400 ml and the day after infected with pGoldenBac virus expressing the six subunits of shelterin. After 60-72 hours cells were collected, washed in PBS 1X and pellets stored at −80°C until further use. Cell pellets were resuspended in 3 volumes of Lysis Buffer (50 mM Tris-HCl pH 7.5, 1 mM DTT, 0.05% Triton-X, 1mM PMSF, 30 µg/ml Leupeptin) supplemented with cOmplete protease inhibitor and PhosSTOP tablets (Merck). After stirring for 15 minutes at 4°C the resuspended pellet was supplemented with 0.5 volumes of glycerol 50%, NaCl concentration adjusted to 300 mM, and stirred for additional 30 minutes at 4°C. Subsequently the lysate was spun down at 48000 g for 45 minutes at 4°C using a JA25.50 rotor. The supernatant was then incubated with 5 ml bed volume Amylose resin (NEB, E8021L) pre equilibrated in MBP Wash buffer (50 mM HEPES pH 7.5, 300 mM NaCl, 1mM Beta-mercaptoethanol, 10% glycerol, 1 mM PMSF). After 1.5 hours shaking at 4°C beads were washed four times with 5 CV of MBP Wash buffer and shelterin eluted in 2 ml fractions using MBP Elution Buffer (50 mM HEPES pH 7.5, 150 mM NaCl, 1mM Beta-mercaptoethanol, 10% glycerol, 1 mM PMSF, 10 mM Maltose). Fractions containing shelterin were pooled and digested overnight with 1:30 w/w PreScission protease to cleave the MBP tags. Shelterin was then loaded on a Hi-Trap Q column (Cytiva) equilibrated in Buffer A (50 mM HEPES pH 7.5, 150 mM NaCl, 10 % glycerol, 1 mM TCEP), washed with 5 CV Buffer A and eluted with a 0-100 % linear gradient of Buffer B (50 mM HEPES pH 7.5, 1 M NaCl, 10 % glycerol, 1 mM TCEP). Fractions containing shelterin were pooled and loaded on a Superose 6 10/300 (Cytiva) and run in SEC buffer (50 mM HEPES pH 7.5, 300 mM NaCl, 10 % glycerol, 1 mM TCEP). Fractions containing shelterin were pooled, concentrated using a Vivaspin 500 (3 kDa) concentrator, aliquoted, snap frozen in liquid nitrogen and stored at −80°C until further use.

Importantly, benzonase was omitted from the purification to avoid potential degradation of individual DNA molecules in single-molecule assays. High salt washes were used to minimise bound genomic DNA. The absence of contaminating DNA following protein purification was first confirmed by the expected A260/A280 absorbance ratio of ∼0.6 for pure protein following size exclusion chromatography (Figure 1B). To further verify the absence of contaminating DNA we also performed AFM experiments with shelterin, TRF1 and TRF2 protein alone (Figure S1C and S1D). No DNA was observed in the protein only images, confirming the cleanliness of the protein preparations (Figure S1C and S1D).

TRF1, TRF2 and associated mutants were purified from insect Sf9 cells. Cells were seeded at 1×10^6^ cells/ml in 400 ml and the day after infected with virus expressing the protein of interest with a HIS tag at its N terminus. After 60-72 hours cells were collected, washed in PBS 1X and pellets stored at −80°C until further use. Cell pellets were resuspended in 3 volumes of Lysis Buffer (50 mM Tris-HCl pH 7.5, 1 mM DTT, 0.1% Nonidet P-40, 1mM PMSF, 30 µg/ml Leupeptin, 20 mM imidazole, supplemented with cOmplete protease inhibitor tablet). After stirring for 15 minutes at 4°C the resuspended pellet was supplemented with 0.5 volumes of glycerol 50%, NaCl concentration adjusted to 300 mM, and stirred for additional 30 minutes at 4°C. Subsequently the lysate was spun down at 48000 g for 45 minutes at 4°C using a JA25.50 rotor. The supernatant was then incubated for 1 hours at 4°C with 1 ml bed volume Ni-NTA agarose (Qiagen) pre equilibrated in Ni-NTA Wash Buffer 1 (50 mM HEPES, pH 7.5, 1M NaCl, 1mM TCEP, 10% glycerol, 40 mM imidazole, 0.1% Nonidet P-40). Beads were then washed twice with 15 CV Ni-NTA wash buffer 1 and an additional four times with 5 CV Ni-NTA Wash Buffer 1 and two times with 5 CV Ni-NTA Wash Buffer 2 (50 mM HEPES, pH 7.5, 300 mM NaCl, 1mM TCEP, 10% glycerol). Elution was performed stepwise with 1 ml of Elution Buffer (50 mM HEPES, pH 7.5, 300 mM NaCl, 1mM TCEP, 10% glycerol, 300 mM imidazole) over 10 fractions. Fractions containing highest concentration of the protein were then dialysed overnight in Dialysis Buffer (50 mM HEPES, pH 7.5, 300 mM NaCl, 1mM TCEP, 10% glycerol) using a Slide-A-Lyzer cassette (Thermo Fisher). After dialysis the protein was collected, aliquoted, snap frozen in liquid nitrogen and stored at −80°C until further use.

### Protein Labelling

For shelterin complex, labelling was performed at a shelterin concentration of 1 µM (ybbr-tag concentration of 10 µM) with 10 mM MgCl₂, 30 uM Alexa-647-CoA and 1 µM Sfp phosphopantetheinyl transferase. Reaction was incubated overnight at 4 °C with gentle rotation. Free dye was removed by size-exclusion chromatography (SEC) using Superose 6 increase 10/300 GL with SEC buffer (50 mM HEPES pH 7.5, 300 mM NaCl, 10 % Glycerol, 1 mM DTT). Fractions were checked on 7% Tris-Acetate Gel and correct fractions pooled and concentrated to ∼1 µM (based on MW of 550 kDa), aliquoted, snap frozen in liquid nitrogen and stored at −80 °C. Final labelled complex was verified, and labelling specificity confirmed, by PAGE gel visualised by fluorescent imaging and Coomassie staining. Labelling efficiency of the individual subunits was estimated to be >90% from absorbance measurements and from the shift of the protein bands observed by PAGE following labelling (Figure S2A).

For TRF1, TRF2 and mutants, labelling was performed at a concentration of 10 µM with 10 mM MgCl₂, 30 uM Alexa-647-CoA and 1 uM Sfp phosphopantetheinyl transferase in 200 µL reactions. Reaction was incubated overnight at 4 °C with gentle rotation. Free dye was removed using 40 kDa Zeba Spin Column and protein aliquoted, snap frozen in liquid nitrogen and stored at −80 °C. Protein and dye concentrations were measured spectrophotometrically. Ratio of protein to dye confirmed labelling efficiencies of 80-100% as previously reported^55,56^. High efficiency labelling was also directly observed by band shift in PAGE gels (Figure S4A).

### Mass photometry

Mass photometry was performed on a Refeyn TwoMP instrument. Imaging slides were constructed using Refeyn MassGlass UC slides (MGUC) and Refeyn Sample Well Cassettes (RD501078). Before assembly, glass slides were washed thoroughly with water and isopropanol. Before adding the sample 10 ul of reaction buffer (50 mM Tris-HCl pH8, 100 mM KCl) was added to the sample well and the correct focus found using autofocus. 10 ul of diluted sample was added to the existing 10 ul droplet before taking a 60 s measurement. Samples were diluted until the number of counts obtained fell within the optimal range as determined by the acquisition software (∼1000 – 3000 counts per minutes). Mass calibration was performed with MassFerence™ P1 (MP-CON-41033) to generate a standard curve and used to convert measured contrast to molecular weight. For TRF2ΔTRFH, whose mass lies close to the lower detection limit of the instrument, measurements were repeated using a reduced field of view and additional slide-cleaning steps to lower the background and improve the limit of detection.

### Gel shift assays

Short double-strand DNA substrates were generated by annealing complementary oligos (Table S1). The substrates used were as follows (with oligo names in parentheses): dsT4 (DD029 and MN1139), dsT4ssT4 (DD029 and MN1137), dsT4ssT2 (DD029 and DD031), dsT2ssT4 (DD030 and DD031) and dsN4 (MN1140 and MN1143). The oligos of 40-nt or longer were first purified on 10% polyacrylamide-urea gels. Oligos were subsequently mixed in equimolar concentrations (10 μM) in STE (100 mM NaCl, 10 mM Tris-HCl at pH 8, 1 mM EDTA), heated, and slowly cooled on a PCR thermocycle (3 min at 98°C, 1 h at 75°C, 1 h at 65°C, 30 min at 37°C, and 10 min at 25°C). For fluorescently labelled dsT4 and dsT4ssT4 substrates, oligo DD029 was replaced by 3′-Atto550 modified version (MN1136). Labelled and unlabelled substrates were purified by native polyacrylamide gel electrophoresis. Binding reactions (20 μL) were carried out in 25 mM Tris-HCl (pH 7.5), 7.5% glycerol, 100 mM NaCl, 2 mM DTT, 1 mM EDTA, and 1 mg/mL BSA. Unless stated otherwise, reactions contained 10 nM fluorescently labelled substrate and the indicated concentration of protein. Shelterin concentrations are expressed as 2:2:2:1:1:1 (TRF1:TRF2:RAP1:TIN2:TPP1:POT1) complex. Complexes were assembled for 30 min at 37°C and separated by gel electrophoresis. Binding reactions were separated on 8% TAE-polyacrylamide gels at 150 V for 2.5 h, and fluorescent gels were visualized using a ChemiDoc system.

### Western blotting of shelterin

Purified shelterin were separated by SDS-PAGE using 4-12% Bis-Tris protein gels and analysed by western blotting. The following primary antibodies were used: rabbit polyclonal anti-POT1(1/500, Novus biological nb500-176), mouse monoclonal anti-TRF1/TRF2 (1/1000, Santa Cruz sc-56807), rabbit polyclonal anti-RAP1 (1/10000, Thermo-Scientific a300-306A), mouse monoclonal anti-TPP1(1/500, Abnova H00065057-MO2) and mouse monoclonal anti-TIN2 (1/2000, Sigma-Aldrich SAB4200108). Secondary antibodies were used at 1/5000: goat anti-mouse IgG (P0447) and swine anti-rabbit IgG (P0399). Western blots were revealed using ChemiDoc system.

### DNA preparation for optical-tweezers and AFM

For optical-tweezers experiments, an 80 telomeric repeat sequence (two blocks of 40 repeats separated by 25 bps) or shorter telomeric repeat sequences were synthesised and cloned into the pMS vector. For AFM analysis, sequences were cloned into the minimal pICOz vector. Plasmids were grown in Stbl3 (Tel_80_) or DH5alpha E. coli strains and extracted using Qiagen MaxiPrep Kits. For AFM, plasmids were further purified by electro-elution. Plasmids were run on 1% TBE agarose gels and the correct band excised and transferred to 10 kDa dialysis tubing (SnakeSkin™, 10K MWCO, 16 mm dry I.D.) with 0.1X TBE buffer. Plasmids were extracted from the gel with electrophoresis for 1 hr at 100v in 0.1X TBE buffer, passed through 0.2 µm spin filter, purified by ethanol precipitation, resuspended in milliQ water and checked by AFM. If required, an additional phenol-chloroform clean-up was performed. Other plasmids used in this work, Tel_12_ and Tel_5_, were prepared in the same manner. The pMS-based plasmids used for optical-tweezers were: Tel_80_ = 3342 bp, Tel_12_ = 2605, Tel_5_ = 2563 bp. The pICOz-based plasmids used for AFM were: Tel_80_ = 1923 bp, Tel_12_ = 1356 bp, Tel_5_ = 1268 bp (Tel_5_), pICOz = 1277 bp.

To generate plasmid arrays for tethering in optical-tweezers, pMS vector plasmids were digested with BstXI. Digested plasmids were diluted to 250 ng/µl in 20 µL of 1x T4 Ligase Buffer and incubated at room temperature for 5 minutes before addition of 1 µL T4 DNA Ligase (NEB, M0202S) for 1 minute followed by addition of first phosphorylated end-cap (End-cap #1, 2 µL, 100 µM, Table S1) for 5 minutes then second phosphorylated end-cap (End-cap #2, 4 uL 100 µM, Table S1) for 5 minutes. Reaction was quenched with 1.5 µL 0.5M EDTA. Ligation was confirmed by running un-ligated and ligated samples on 0.5% TAE agarose gel at 100 V for 1 hr.

### Optical-tweezers experiments

The optical-tweezers experiments were performed on a commercial LUMICKS C-trap optical-tweezers microscope with integrated confocal microscopy and microfluidics (Serial Number BA105). The flow cell and fluidics tubing was cleaned with 1 ml 2% Hellmanex™ III (Z805939) in each syringe, flowed over 30 minutes, followed by 1 ml MQ H_2_O and 0.5 ml wash buffer (50 mM Tris-HCl pH8, 100 mM KCl). Protein channels were passivated with > 500 µL wash buffer supplemented with 0.1% Casein over 30 minutes then washed with 500 µL reaction buffer (50 mM Tris-HCl pH8, 100 mM KCl, 0.001% Tween-20) before addition of proteins. Beads were diluted to 0.005% w/v and DNA diluted 1:200 in reaction buffer. The ligated telomeric plasmid arrays were tethered between optical-trapped polystyrene particles (4.38 µm streptavidin-coated SPHERO) using the microfluidic flow cell. Force-distance curves were performed in reaction buffer to verify the presence of a single DNA molecule and to calculate the DNA contour length. Proteins were diluted in reaction buffer (50 mM Tris-HCl pH8, 100 mM KCl, 1 mM DTT, 0.001% Tween-20). DNA was held at 5 pN before being moved into the protein channel and imaged.

2D scans were performed with 50 nm pixel size, 0.1 ms/pixel dwell time and 2% red laser power (647 nm). Kymographs were generated with 50 nm pixel size, 0.1 ms/pixel dwell time, 100 ms line time. Images were exported as .tiff files using Pylake python package and processed in Fiji.

The distance between the beads was then reduced to 5 µm and incubating for 10 s before performing force-distance curves from 5 µm distance to 55 pN with a constant pulling speed of 1 µm/second. Force-distance analysis was performed in Python using pylake and custom analysis scripts which have been deposited with the data. Rupture forces were determined using peak finding from force-time traces, rupture forces were extracted from three cycles of pulling and relaxation per DNA molecule and at least three different DNA molecules were measured for each condition. The unit of replication is different individual DNA molecules, not the individual rupture events. DNA arrays were prepared fresh and used for a maximum of 2 weeks, protein preparations used were prepared once. Statistical significance was determined using Kruskal–Wallis test followed by post-hoc Dunn test with Holm correction, * = p ≤ 0.05, ** = p ≤ 0.01 and *** = p ≤ 0.001. Force-distance plotting, eWLC fitting and contour length transformations were performed in Igor Pro 9 (Version 9.01). Contour length transformations were preformed using Equation 2. with fixed persistence length, Lp = 50 nm and stretch modulus, S_t_ = 1500 pN.

For protein dissociation measurements, tethered DNA arrays were held at 5 pN and incubated in the protein channel with 20 nM labelled TRF1 or TRF2 for 1 minute before being moved to a buffer channel containing no protein, and kymographs were recorded for 300 s. Analysis was performed in Python using pylake and custom analysis scripts which have been deposited with the data. Briefly, binding sites were identified across region corresponding to the DNA by peak finding of summed intensity profile in x. Intensity-time traces were extracted for individual telomeric binding sites and fitted with a single exponential decay to give the dissociation rate (1/τ); the initial bound intensity was taken as the fitted intensity at t = 0. Statistical significance was determined using Kruskal–Wallis test followed by post-hoc Dunn test with Holm correction, * = p ≤ 0.05, ** = p ≤ 0.01 and *** = p ≤ 0.001.

For trans capture experiments double-stranded DNA substrates were generated by annealing equimolar ratios of oligos MN1136 + MN1139 (Table S1) for the telomeric repeat substrate (ds4T) and MN1140 + MN1143 (Table S1) for the scrambled, non-telomeric repeat substrate (ds4N). Tethered DNA was held a 10 pN force before moving to position in the flow cell containing 2 nM annealed oligo and 10 nM unlabelled protein in reaction buffer. For analysis of capture, horizontal line intensities were measured for 10 µm region (200 pixels) of the kymograph corresponding to the DNA and normalised by time. Line intensities were measured covering at least 10 binding sites per DNA molecule from at least three kymographs from different DNA molecules. Statistical significance between groups was determined using Kruskal–Wallis test followed by post-hoc Dunn test with Holm correction, * = p ≤ 0.05, ** = p ≤ 0.01 and *** = p ≤ 0.001.

### AFM sample preparation

DNA sample preparation was conducted following a published protocol^57,58^. Sample preparation for AFM imaging of DNA samples was performed by depositing ∼5 ng of DNA into a 30 µL droplet of immobilisation buffer (3 mM NiCl_2_, 20 mM HEPES, pH7.4) on discs of freshly cleaved muscovite mica. The sample disc was incubated for 5 minutes before washing three times with fresh immobilisation buffer to remove any non-adsorbed DNA, before adding 30 µL of fresh buffer for imaging. For imaging samples with protein, each protein was diluted to 20 nM in dilution buffer (50 mM tris, 2 mM MgCl_2_ 5% glycerol) and then co-incubated with DNA and incubation buffer (100 mM TRIS, 4 mM MgCl_2_, 200 mM KCl, 10% glycerol, pH 8) to a final concentration of 1 ng/µL DNA and 5 nM protein. Unless stated otherwise, all AFM experiments were performed at 5 nM protein; for the higher-concentration experiments shown in Figures S3G–S3J, proteins were added to final concentrations of 10 nM and 20 nM respectively, with all other conditions unchanged. The sample was then immobilised on the mica as before. The unit of replication is different individual complexes, with each taken as an individual image. The same protein preparations were used across the optical-tweezers and AFM experiments. A single preparation of each DNA plasmid was used across all samples/conditions. To ensure any variation was due to protein behaviour rather than DNA variability. Samples were complexed fresh at the start of each imaging session, each DNA:protein complex was prepared at least twice.

### AFM imaging

Sample scans were captured using a Dimension XR FastScan (Bruker) using FastScanD cantilevers (Bruker) and PeakForce Tapping mode. The PeakForce amplitude was set to 6-10 nm, the PeakForce Tapping frequency to 8 kHz and the PeakForce setpoints in the range: 7–15 mV, corresponding to peak forces of <70 pN. The number of pixels recorded for each image was adjusted to maintain a resolution of ≤0.5 nm/pixel.

### AFM image analysis

AFM images were processed using the open-source AFM image analysis software TopoStats (https://github.com/AFM-SPM/TopoStats, v2.4.2)^59,60^. The full configuration file used can be found in the SI and full documentation can be found in the TopoStats documentation (information can be found here https://github.com/AFM-SPM/TopoStats and the configuration file used is included in the data set).

Flattening was performed using TopoStats’ default settings, where row alignment and then tilt removal eliminated scan lines and flattened the background, then the image’s background height was zeroed. A second round of filtering was then performed, ignoring regions greater than 1 standard deviation away from the image’s background as to prevent DNA data from affecting the flattening process. Finally, a 1px gaussian blur was applied to reduce the effects of high frequency noise.

Grains of interest were identified using an absolute height threshold of 1.0 nm and a minimum area of 1000 nm^2^. Grains intersecting the image border were removed and remaining grains were then passed to a U-Net model for multi-class re-segmentation^57,60^. A U-Net deep learning segmentation model was trained using 70 hand labelled images, each undergoing a set of 5 augmentations including a combination of horizontal and vertical flips and random cropping, which increased the training set to a total of 420 paired AFM images and masks. The U-Net was trained for 200 epochs and used a dice coefficient loss function that ignored background class. Two classes were used for the labels, one marking regions of DNA, and the other marking regions of protein, designed to enable more powerful analysis. Validation set accuracy for this model at the final epoch was 0.98. The resulting U-Net model was used to re-segment grains of interest, resulting in more successful downstream processing and protein identification.

Basic morphological statistics were then calculated for the grains, such as height, area, volume, and aspect ratio, for both the DNA and protein classes, before being passed to TopoStats’ DNA tracing pipeline. The DNA tracing pipeline identified crossing points and discerned different DNA branches and calculated their lengths. Statistical significance was determined using Kruskal–Wallis test followed by post-hoc Dunn test with Holm correction, * = p ≤ 0.05, ** = p ≤ 0.01 and *** = p ≤ 0.001.

For comparison of protein mediated plasmid tethering between conditions, we calculated a normalised tethering factor. This metric was calculated by first normalising the DNA length and protein volume values between 0 and 1 across all molecules in the dataset, pooling every condition being compared, so that the resulting values remain directly comparable between conditions. The tethering factor is then a multiplication of the normalised DNA length and normalised protein volume for each complex. This combined DNA length and protein volume metric is required to overcome the technical limitation of AFM that DNA can become hidden on protein binding resulting in metrics which only account for changes in DNA length not accurately describing the data.

Using our automated pipeline, we determine the variation across the data set due to tip variation, calculated from the observed DNA width, and changes in imaging parameters, calculated from measurements of DNA height, i.e. molecular compression, across sessions to allow for comparison across samples (Figure S5). Calculating the DNA width and height across samples and datasets we observe ∼10% variability (CV) in DNA deformation (i.e. height) (Figure 5SA) and ∼20% variability (CV) in tip convolution (Figure S5B).

### Fluorescence polarisation

For fluorescence polarisation assays protein was buffer exchanged using Zeba™Spin Desalting Columns (7K MWCO, 0.5 mL, 89882) into 50mM Tris pH8.0, 100mM KCl, with 1/1000x protease inhibitor cocktail (Roche P8340). The protein concentration was measured (Bradford) and diluted to 2 uM. Different concentrations of protein were incubated for 15 mins with 70 pM pre-annealed FAM-labelled oligonucleotides containing 4 x telomeric repeats (MN2043 + MN2044, Table S1) in a 384 well plate (Corning® Low Volume 384-well Black/Clear Flat Bottom Polystyrene Not Treated Microplate, 3540). Non-telomeric DNA control assays were performed as above but using oligonucleotides with scrambled telomeric repeats (MN2045 + MN2046, Table S1). Fluorescence polarisation was measured on a Hidex sense plate reader using 200 flashes of 485/10nm excitation, 520/14nm emission, with a 505nm dichromic mirror. Specific affinity for binding to telomeric DNA was calculated by subtracting mean mP for non-telomeric DNA from that of telomeric DNA (n ≥ 3). Data was fit with Hill equation.

### Crosslinking Mass Spec

For crosslinking Mass Spec 50 pmol of shelterin were crosslinked with 100X molar excess of DSBU (Thermo Fisher) for 2 hours at 4°C. After crosslinking samples were quenched with 100 mM Tris-HCl pH 7.5 for 30 minutes at room temperature.

Crosslinked complexes were reduced with 10 mM dithiothreitol and alkylated with 50 mM iodoacetamide. They were then digested with trypsin at an enzyme-to-substrate ratio of 1:100, for 1 hr at RT and further digested overnight at 37°C following addition of trypsin at a ratio of 1:20. The peptide digests were then fractionated batch-wise by high pH reverse phase chromatography on micro spin TARGA C18 columns (Nest Group Inc, USA), into five fractions (10 mM NH4HCO3/10% (v/v) acetonitrile pH 8.0, 10 mM NH4HCO3/20% (v/v) acetonitrile pH 8.0, 10 mM NH4HCO3/30% (v/v) acetonitrile pH 8.0, 10 mM NH4HCO3/40% (v/v) acetonitrile pH 8.0 and 10 mM NH4HCO3/80% (v/v) acetonitrile pH 8.0). The fractions (120 µL) were evaporated to dryness in a CentriVap concentrator (Labconco, USA) prior to analysis by LC-MS/MS.

Lyophilized peptides were resuspended in 1% (v/v) formic acid and 2% (v/v) acetonitrile and analysed by nano-scale capillary LC-MS/MS using a Vanquish Neo UPLC (ThermoScientific Dionex, USA) to deliver a flow of approximately 250 nL/min. A PepMap Neo C18 5 μm, 300 μm x 5 mm nanoViper (ThermoScientific Dionex, USA) trapped the peptides before separation on a 50 cm EASY-Spray column (50 cm x 75 µm ID, PepMap C18, 2 µm particles, 100 Å pore size: ThermoScientific, USA). Peptides were eluted with a gradient of acetonitrile over 90 minutes. The analytical column outlet was directly interfaced via a nano-flow electrospray ionisation source, with a quadrupole Orbitrap mass spectrometer (Orbitrap Exploris 480, ThermoScientific, USA). MS data were acquired in data-dependent mode using a top 10 method, where ions with a precursor charge state of 1+ and 2+ were excluded. High-resolution full scans (R=60000, m/z 380-1800) were recorded in the Orbitrap followed by higher energy collision dissociation (HCD) (stepped collision energy 30 and 32% Normalized Collision Energy) of the 10 most intense MS peaks. The fragment ion spectra were acquired at a resolution of 15,000 and a dynamic exclusion window of 20 secs was applied.

For data analysis, Xcalibur raw files were converted into the MGF format using Proteowizard^61^ and used directly as input files for MeroX^62^. Searches were performed against an ad hoc protein database containing the sequences of the proteins in the complex and a set of randomized decoy sequences generated by the software. The following parameters were set for the searches: maximum number of missed cleavages: 3; targeted residues K, S, Y and T; minimum peptide length 5 amino acids; variable modifications: carbamidomethylation of cysteine (mass shift 57.02146 Da), methionine oxidation (mass shift 15.99491 Da); DSBU modified fragments: 85.05276 Da and 111.03203 Da (precision: 5 ppm MS and 10 ppm MS/MS); False Discovery Rate cut-off: 5%. Resulting Cross-links were visualized in xiView^63^.

## Acknowledgements

This work was supported by a UKRI Future Leaders Fellowship MR/W00738X/1 to A.L.B.P. and Wellcome Early Career Award (225139/Z/22/Z) to M.D.N., MRC Equipment Grant (MC PC APP28599) for LUMICKS C-trap Instrument, Wellcome Trust and the Royal Society Grant (213501/Z/18/Z) for plate reader experiments and BBSRC ALERT Mid-range equipment award (BB/Z515930/1) for mass photometry experiments. Work in the S.J.B. lab is supported by the Francis Crick Institute (CC2098), ERC Advanced Investigator and UKRI grants (TelMetab, ChrEndProt), a Wellcome Trust Senior Investigator Award, and CRUK RadNet City of London. L.G. is supported by the European Union’s Horizon research and innovation programme under the Marie Sklodowska-Curie grant agreement no. 101203604. We wish to acknowledge the Henry Royce Institute for Advanced Materials, funded through EPSRC grants EP/R00661X/1, EP/S019367/1, EP/P02470X/1 and EP/P025285/1 and Xinyue Chen for Dimension FastScan access and support at Royce@Sheffield.

## Author Contributions

Formal contributions in authorship order (CrediT taxonomy): Conceptualization: M.D.N., A.L.B.P., S.J.B.; Data curation: M.D.N., T.E.C., S.W., L.W., E.P.H., E.G.A.; Formal Analysis: M.D.N., T.E.C., S.W., L.W., E.G.A., S.M.; Funding acquisition: M.D.N., A.L.B.P., S.J.B.; Investigation: M.D.N., T.E.C., L.G., D.D., E.P.H., E.G.A., M.G., S.M.; Methodology: M.D.N., S.W., L.G., L.W., E.P.H., D.D., R.A., E.G.A., S.S.-B., O.B., M.C.G., M.S., A.L.B.P. and S.J.B.; Project administration: M.D.N., A.L.B.P., and S.J.B.; Resources: M.D.N., A.L.B.P. and S.J.B.; Software: S.W., L.W., M.C.G.; Supervision: M.D.N., A.L.B.P. and S.J.B; Visualization: M.D.N., T.E.C.; Writing – original draft: M.D.N, T.E.C., A.L.B.P. and S.J.B.; Writing – review & editing: M.D.N., T.E.C., S.W., L.W., E.P.H, A.L.B.P. and S.J.B.

## Data Availability

The mass spectrometry proteomics data have been deposited to the ProteomeXchange Consortium via the PRIDE^64^ partner repository (https://www.ebi.ac.uk/pride/) with the dataset identifier PXD077320. The raw and processed AFM data can be found on FigShare at the following DOI: 10.15131/shef.data.33269223. The optical-tweezers data can be found on Figshare DOI: 10.6084/m9.figshare.33254085). Source Data are provided with this paper.

## Code Availability

All AFM analysis code is publicly available on GitHub via TopoStats (https://github.com/AFM-SPM/TopoStats, v2.4.2). The configuration file for TopoStats, the U-Net model and the python notebook used to compute the normalised tethering factor can be found on FigShare at the following DOI: 10.15131/shef.data.33269223.

The python notebooks used for optical-tweezers data analysis and plotting can be found on FigShare with the corresponding raw data at the following DOI: 10.6084/m9.figshare.33254085.

## Supplementary Information

Supplementary information is available containing Supplementary Figures S1 to S6 and Supplementary Tables S1 and S2.

