## Supplementary Information for "TRF2 mediates force-resistant bridging of distal telomeric DNA repeats"

**The supplementary information file includes:**

Supplementary Figures S1 to S6

Supplementary Tables S1 and S2

### Supplementary Figures

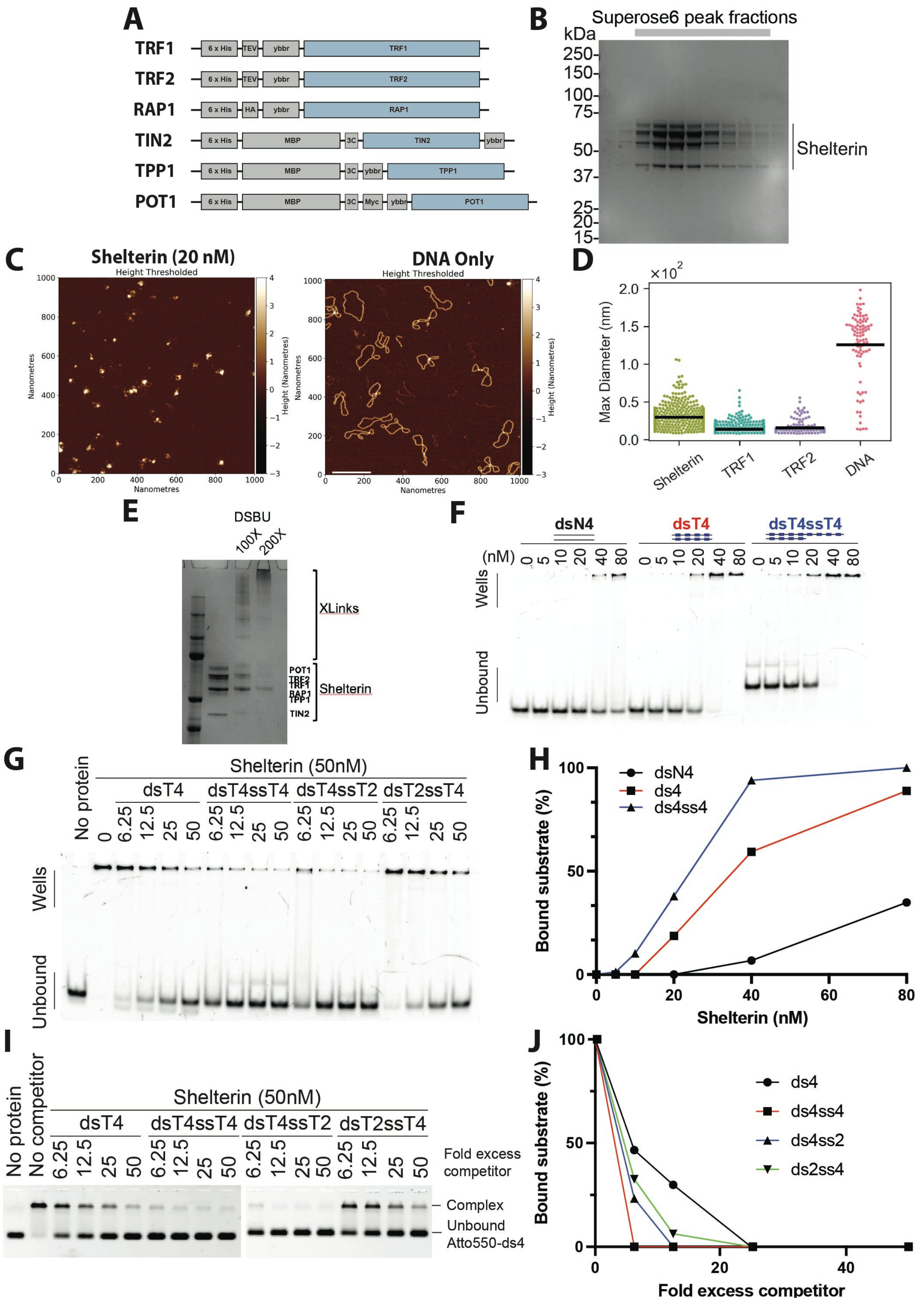

**Figure S1. Reconstituted shelterin complexes contain both TRF1 and TRF2 and are active in telomeric DNA binding.** (A) Schematic of the GoldenBac system for co-expression of all six shelterin proteins from a single bacmid. (B) SDS-PAGE gel of Superose6 peaks. Fractions from grey shaded region in Figure 1B highlighted with top grey bar. (C) Representative AFM image of Shelterin protein prep (Scale bar = 200 nm), Left. Representative AFM image of pICOz plasmid DNA Only (1.27 kb), Right (D) Quantification of diameter of detected particles. Shelterin, n = 526; TRF1, n = 765; TRF2, n = 141; pICOz plasmid DNA, n = 87. (E) SDS-PAGE gel of shelterin DSBU crosslinking at 100x and 200x. (F) Example polyacrylamide gel EMSA of purified shelterin binding to different telomeric and non-telomeric DNA substrates. dsN4 = 4 x double-stranded scrambled telomeric repeats, dsT4 = 4 x double-stranded telomeric repeats, dsT4ssT4 = 4 x double-stranded telomeric repeats with 4 x single-stranded telomeric repeat overhang. (G) Example competition polyacrylamide gel EMSA of purified shelterin bound to four double-stranded telomeric repeats competed with different telomeric substrates. (H) Quantification of agarose gel EMSA of purified shelterin binding to different telomeric and non-telomeric DNA substrates, Figure 1G. dsN4 = 4 x double-stranded scrambled telomeric repeats, dsT4 = 4 x double-stranded telomeric repeats, dsTT4 = 4 x double-stranded telomeric repeats with 4 x single-stranded telomeric repeat overhang. (I-J) Agarose gel competition EMSA (I), and quantification (J), of purified shelterin bound to four double-stranded telomeric repeats competed with telomeric DNA substrates containing different numbers of double-stranded and single-stranded telomeric repeats.

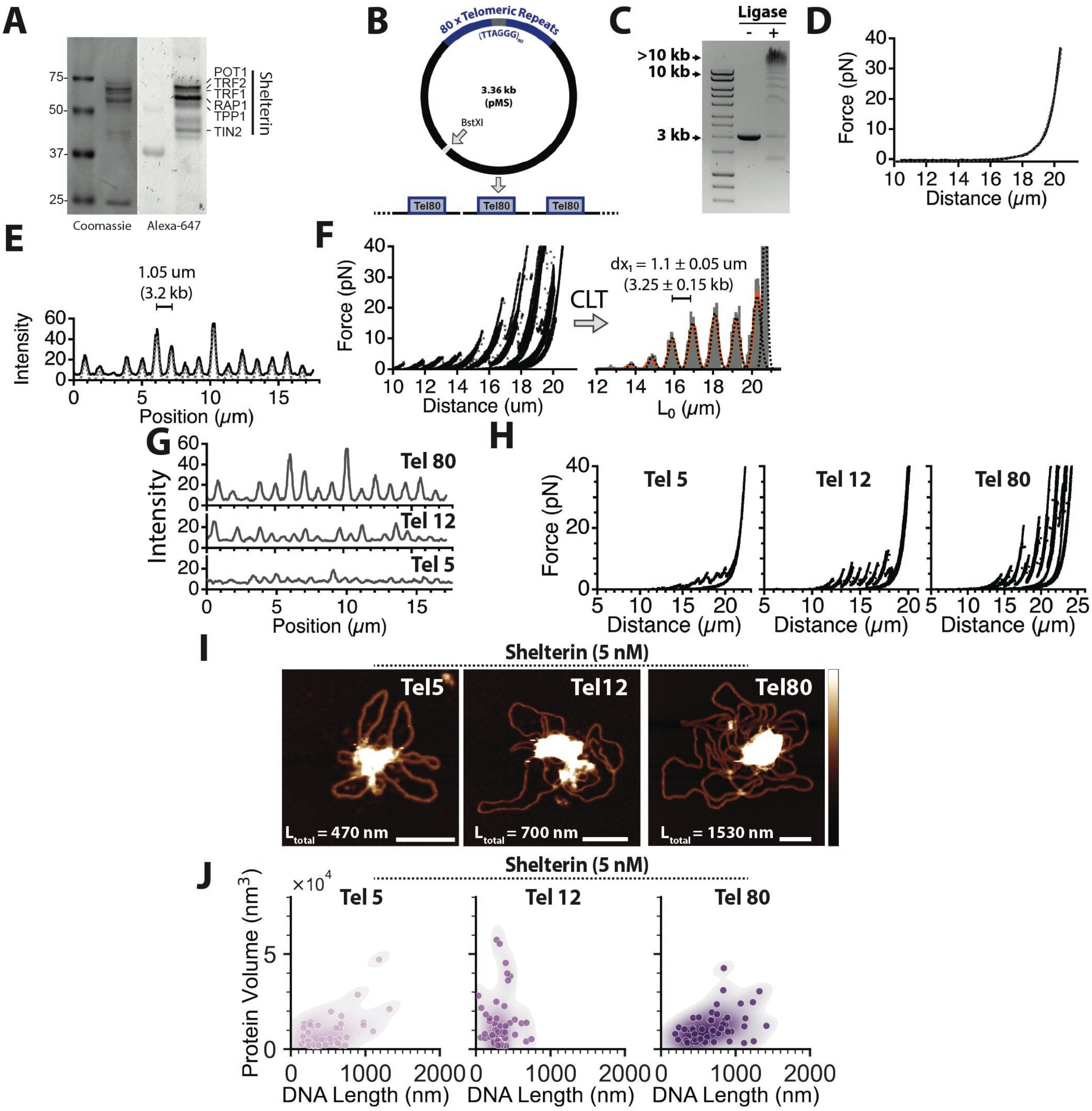


**Figure S2. Shelterin mediates tethering between telomeric repeats.** (A) SDS-page gel of purified Alexa-647 labelled shelterin showing specific labelling of shelterin proteins. Left, coomassie stained, right, fluorescence imaging (647 nm) prior to staining. (B) Schematic of generation of long DNA arrays containing blocks of telomeric repeats. (C) Agarose gel electrophoresis of digested plasmid before and after ligation showing generation of high molecular weight DNA, right. (D) Force-distance curve of tethered $Tel_{80}$ DNA array. Fit with extensible worm-like chain model, dashed line. (E) Example intensity profile for $Tel_{80}$ DNA array incubated with labelled shelterin. (F) Force-distance data from repeated pulling and relaxation cycles of $Tel_{80}$ incubated with shelterin, left, contour length transformation (CLT) of force-distance rupture data to calculate distributions of DNA lengths. CLT data peaks fit with gaussian fits to calculate interpeak distances, $dx_1$. (G) Representative fluorescence intensity plots for $Tel_5$, $Tel_{12}$ and $Tel_{80}$ DNA incubated with fluorescently labelled shelterin (H)

Representative force-distance curves of repeated pulling and relaxation cycles of $Tel_5$, $Tel_{12}$ and $Tel_{80}$ DNA arrays incubated with shelterin (I) Representative AFM images of $Tel_5$, $Tel_{12}$ and $Tel_{80}$ plasmids incubated with shelterin (5 nM). Scale bars = 50 nm, z scale = -2 to 5 nm. (J) Quantification of protein volume vs total DNA length for $Tel_5$, $Tel_{12}$ and $Tel_{80}$ plasmids incubated with shelterin (5 nM). N values: $Tel_5$ = 48, $Tel_{12}$ = 60, $Tel_{80}$ = 50. Statistical significance between groups assessed with a post-hoc Dunn's test.

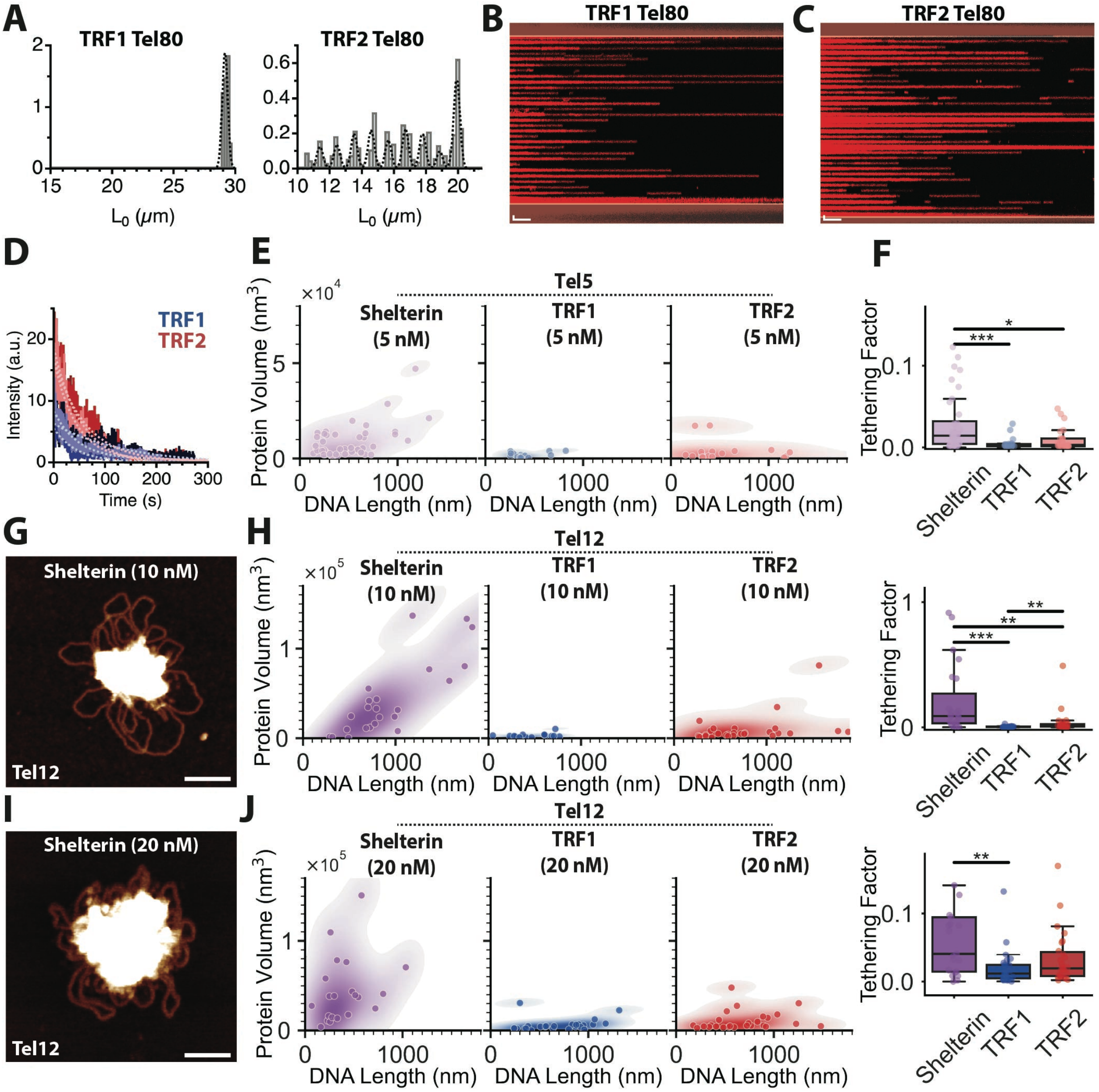


**Figure S3. TRF2 alone is sufficient to mediate telomeric DNA tethering.** (A) Contour length transformation of $Tel_{80}$ TRF1 and TRF2 force-distance curves in Figures 3A and 3B. Fit with single gaussian, TRF1, left, or multi-gaussian, TRF2, right (B) Representative kymograph of dissociation of prebound TRF1 from $Tel_{80}$ array in buffer only channel. Scale bars: x = 10 s, y = 1 µm. (C) Representative kymograph of dissociation of prebound TRF2 from $Tel_{80}$ array in buffer only channel. Scale bars: x = 10 s, y = 1 µm. (D) Representative extracted intensity-time plots of dissociation of TRF1, blue, and TRF2, red, from $Tel_{80}$ arrays. Fit with single exponential decay, grey dashed lines. (E) Quantification of protein volume vs total DNA length observed by AFM for $Tel_5$ plasmid incubated with 5 nM shelterin, TRF1 or TRF2. (F) Comparison of calculated normalized tethering factor from AFM measurements of $Tel_5$ plasmids incubated with 5 nM shelterin, TRF1 or TRF2. N values: shelterin = 48, TRF1 = 18. TRF2 = 23. Statistical significance between groups assessed with a post-hoc Dunn's test. (G) Representative AFM image for $Tel_{12}$ plasmid incubated with shelterin (10 nM). Scale bars =

50 nm, z scale = -2 to 5 nm. (H) Quantification of protein volume vs total DNA length observed by AFM for $Tel_{12}$ telomeric repeat-containing plasmids incubated with 10 nM shelterin, TRF1 and TRF2, left. Corresponding comparison of calculated normalized tethering factor. N values: shelterin = 23, TRF1 = 25, TRF2 = 30. Statistical significance between groups assessed with a post-hoc Dunn's test. (I) Representative AFM image for $Tel_{12}$ plasmid incubated with shelterin (20 nM). Scale bars = 50 nm, z scale = -2 to 5 nm. (J) Quantification of protein volume vs total DNA length observed by AFM for $Tel_{12}$ telomeric repeat-containing plasmids incubated with 20 nM shelterin, TRF1 and TRF2, left. Corresponding comparison of calculated normalized tethering factor. N values: shelterin = 22, TRF1 = 25, TRF2 = 26. Statistical significance between groups assessed with a post-hoc Dunn's test.

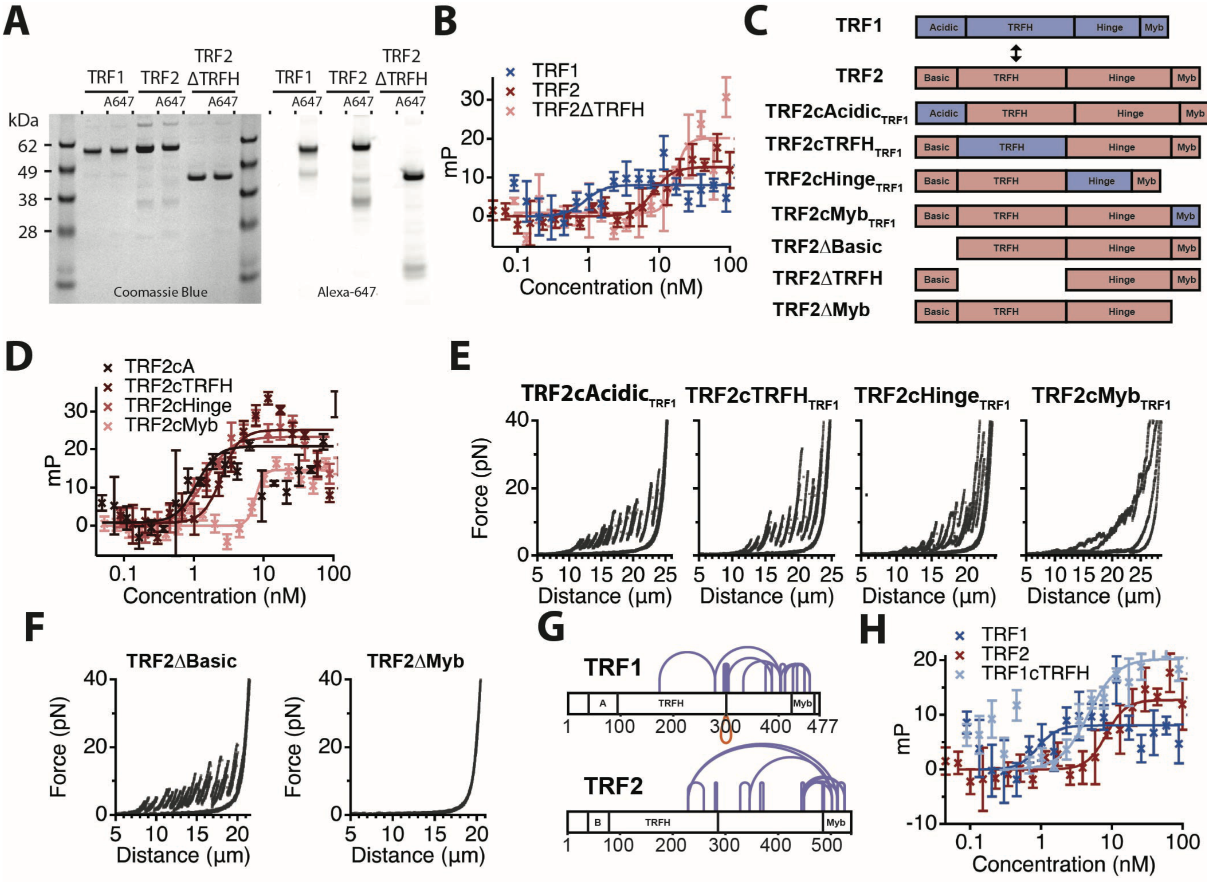


**Figure S4. TRF2 mediated tethering involves multiple domains of TRF2.** (A) SDS-PAGE gel of purified unlabelled and Alexa-647 fluorescently labelled TRF1, TRF2 and TRF2ΔTRFH. Coomassie stained image, left, and fluorescent image (647 nm) prior to staining, right (B) Plot of fluorescence polarisation measurements of specific binding to telomeric repeats for TRF1, TRF2 and TRF2ΔTRFH, n ≥ 3. Fit with Hill Equation. (C) Schematic of domain swap mutants of TRF2. (D) Plot of fluorescence polarisation measurements of specific binding to telomeric repeats for all four TRF2 domain swap mutants, n ≥ 3. Fit with Hill Equation. (E) Representative force-distance curves showing tethering for $Tel_{80}$ DNA array incubated with TRF2 domain swap mutants. (F) Representative force-distance curve for $Tel_{80}$ incubated with TRF2ΔBasic, left, and TRF2ΔMyb, right. (G) Crosslinking mass spectrometry analysis of intra- TRF1 and TRF2 crosslinks from analysis of the purified shelterin complex. (H) Plot of fluorescence polarisation measurements of specific binding to telomeric repeats for TRF1, TRF2 and $TRF1cTRFH_{TRF2}$, n ≥ 3. Fit with Hill Equation.

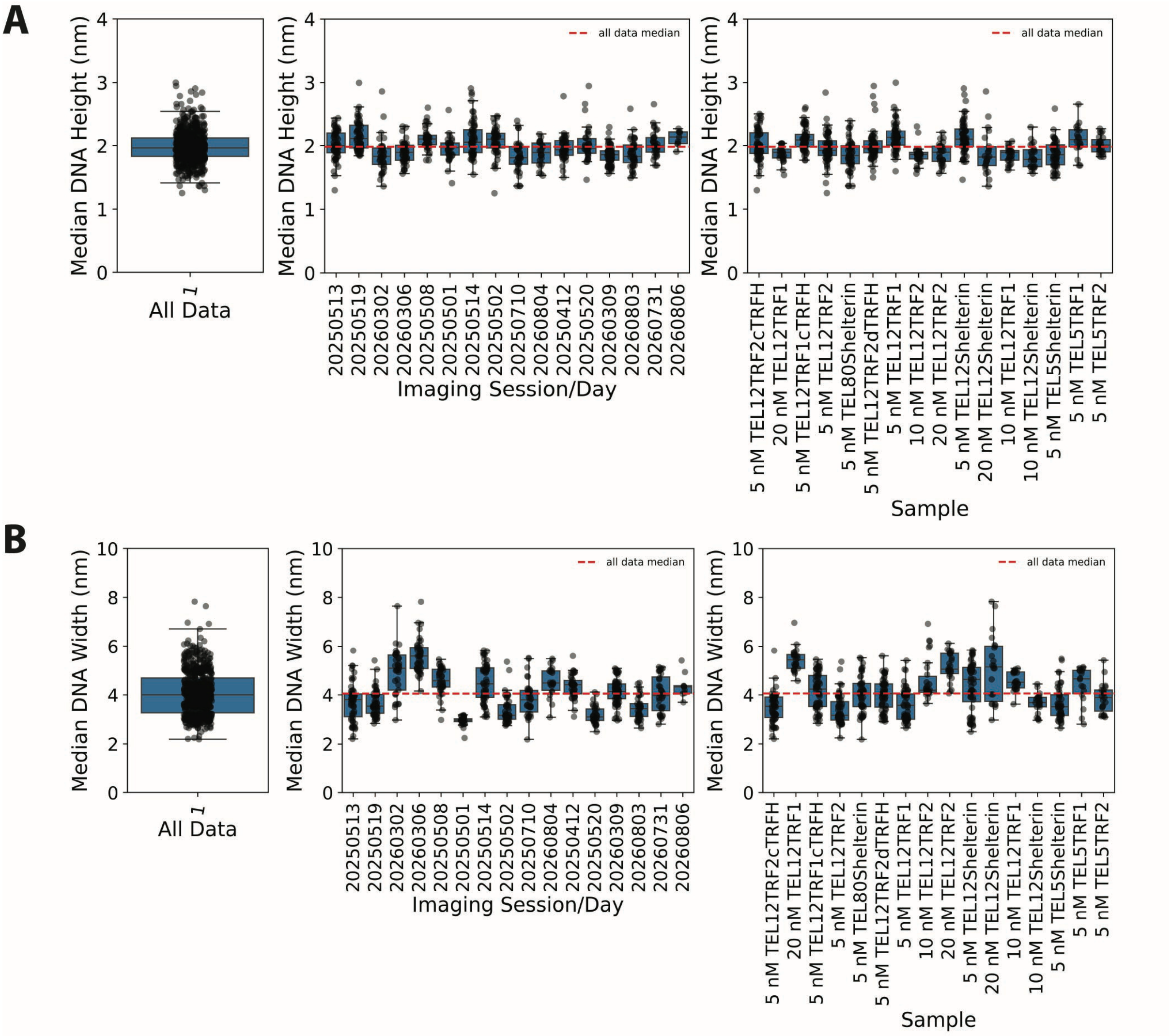


**Figure S5. Comparison of AFM imaging conditions as a function of DNA height (force applied) and DNA width (tip convolution).** (A) Median measured DNA height for each molecule for whole dataset, left, and separated by imaging session/day, middle, or by sample, right. Red dotted line, all data median value. (B) Median measured DNA width for each molecule for whole dataset, left, and separated by imaging session/day, middle, or by sample, right. Red dotted line, all data median value.

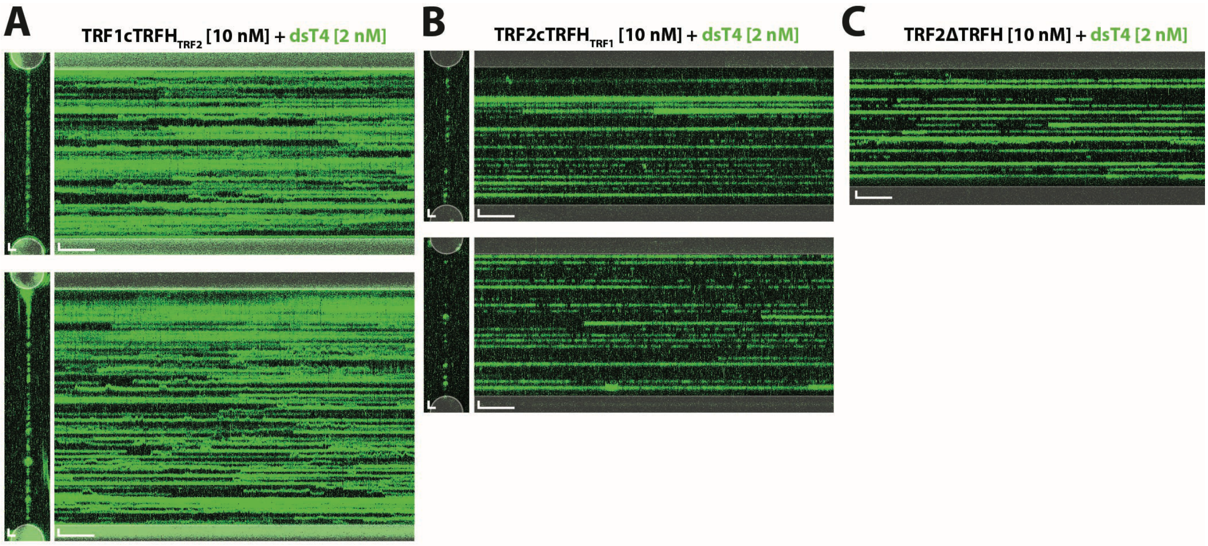


**Figure S6. TRF2 TRFH domain mediates capture of telomeric DNA in trans.** (A) 2D-scans and corresponding kymographs for incubation of $Tel_{12}$ DNA arrays with $TRF1cTRFH_{TRF2}$ (10 nM) and labelled 4xTel dsDNA. 2D-scan scale bars = 1 µm. Kymograph scale bars: x = 10 s, y = 1 µm, (B) 2D-scans and corresponding kymographs for incubation of $Tel_{12}$DNA arrays with $TRF2cTRFH_{TRF1}$ (10 nM) and labelled 4xTel dsDNA. 2D-scan scale bars = 1 µm. Kymograph scale bars: x = 10 s, y = 1 µm. (C) Kymographs for incubation of $Tel_{12}$ DNA arrays with TRF2ΔTRFH (10 nM) and labelled 4xTel dsDNA. Kymograph scale bars: x = 10 s, y = 1 µm.

**Table S1**

| Name | Sequence (5'-3') |
|---|---|
| MN1136 | (CCCTAA)$_4$GCCTCGCTGCCGTCGCCA/Atto550 |
| MN1137 | TGGCGACGGCAGCGAGGC(TTAGGG)$_8$ |
| MN1139 | TGGCGACGGCAGCGAGGC(TTAGGG)$_4$ |
| MN1140 | (CACTCA)$_4$GCCTCGCTGCCGTCGCCA/Atto550 |
| MN1143 | TGGCGACGGCAGCGAGGC(TGAGTG)$_4$ |
| DD029 | (CCCTAA)$_4$GCCTCGCTGCCGTCGCCA |
| DD030 | (CCCTAA)$_2$GCCTCGCTGCCGTCGCCA |
| DD031 | TGGCGACGGCAGCGAGGC(TTAGGG)$_6$ |
| Endcap #1 | GGAGTTGAACG/iBiodT//iBiodT/T/iBiodT/T/iBiodT/ACGTTCAACTCCCACT |
| Endcap #2 | CAAGTTGGACAA/iBiodT/T/iBiodT/T/iBiodT/T/iBiodT/TGTCCAACTTGAGTG |
| MN2043 | tgccagactcTTAGGGTTAGGGTTAGGGTTAGGG |
| MN2044 | CCCTAACCCTAACCCTAACCCTAAgagtctggca/FAM/ |
| MN2045 | tgccagactcTGAGTGTGAGTGTGAGTGTGAGTG |
| MN2046 | CACTCACACTCACACTCACACTCAgagtctggca/FAM/ |

### Table S2

| Name | Amino acid sequence |
|---|---|
| TRF1 | MGHHHHHHGTGSENLYFQGADSLEFIASKLAGSGSGSGMAEDVSSAAPSPRGCADGRDADPTEEQMAETERNDEEQFECQ<br>ELLECQVQVGAPEEEEEEEEDAGLVAEAEAVAAGWMLDFLCLSLCRAFRDGRSEDFRRTRNSAEAIIHGLSSLTACQLRT<br>IYICQFLTRIAAGKTLDAQFENDERITPLESALMIWGSIEKEHDKLHEEIQNLIKIQAIAVCMENGNFKEAEEVFERIFG<br>DPNSHMPFKSKLLMIISQKDTFHSFFQHFSYNHMMEKIKSYVNYVLSEKSSTFLMKAAAKVVESKRTRTITSQDKPSGND<br>VEMETEANLDTRKSVSDKQSAVTESSEGTVSLLRSHKNLFLSKLQHGTQQQDLNKKERRVGTPQSTKKKKESRRATESRI<br>PVSKSQPVTPEKHRARKRQAWLWEEDKNLRSGVRKYGEGNWSKILLHYKFNNRTSVMLKDRWRTMKKLKLISSDSED |
| TRF2 | MGHHHHHHGTGSENLYFQGAGDSLEFIASKLAGSGSGSGMAGGGGSSDGSGRAAGRRASRSSGRARRGRHEPGLGGPAER<br>GAGEARLEEAVNRWVLKFYFHEALRAFRGSRYGDFRQIRDIMQALLVRPLGKEHTVSRLLRVMQCLSRIEEGENLDCSFD<br>MEAELTPLESAINVLEMIKTEFTLTEAVVESSRKLVKEAAVIICIKNKEFEKASKILKKHMSKDPTTQKLRNDLLNIIRE<br>KNLAHPVIQNFSYETFQQKMLRFLESHLDDAEPYLLTMAKKALKSESAASSTGKEDKQPAPGPVEKPPREPARQLRNPPT<br>TIGMMTLKAAFKTLSGAQDSEAAFAKLDQKDLVLPTQALPASPALKNKRPRKDENESSAPADGEGGSELQPKNKRMTISR<br>LVLEEDSQSTEPSAGLNSSQEAASAPPSKPTVLNQPLPGEKNPKVPKGKWNSSNGVEEKETWVEEDELFQVQAPDEDSTT<br>NITKKQKWTVEESEWVKAGVQKYGEGNWAAISKNYPFVNRTAVMIKDRWRTMKRLGMN |
| TRF1cTRFH$_{TRF2}$ | MGHHHHHHGTGSENLYFQGADSLEFIASKLAGSGSGSGMAEDVSSAAPSPRGCADGRDADPTEEQMAETERNDEEQFECQ<br>ELLECQVQVGAPEEEGAGEARLEEAVNRWVLKFYFHEALRAFRGSRYGDFRQIRDIMQALLVRPLGKEHTVSRLLRVMQC<br>LSRIEEGENLDCSFDMEAELTPLESAINVLEMIKTEFTLTEAVVESSRKLVKEAAVIICIKNKEFEKASKILKKHMSKDP<br>TTQKLRNDLLNIIREKNLAHPVIQNFSYETFQQKMLRFLESHLDDAEPYLLTMAKKALKRTRTITSQDKPSGNDVEMETE<br>ANLDTRKSVSDKQSAVTESSEGTVSLLRSHKNLFLSKLQHGTQQQDLNKKERRVGTPQSTKKKKESRRATESRIPVSKSQ<br>PVTPEKHRARKRQAWLWEEDKNLRSGVRKYGEGNWSKILLHYKFNNRTSVMLKDRWRTMKKLKLISSDSED |
| TRF2cTRFH$_{TRF1}$ | MGHHHHHHGTGSENLYFQGAGDSLEFIASKLAGSGSGSGMAGGGGSSDGSGRAAGRRASRSSGRARRGRHEPGLGGPAER<br>EEEEEDAGLVAEAEAVAAGWMLDFLCLSLCRAFRDGRSEDFRRTRNSAEAIIHGLSSLTACQLRTIYICQFLTRIAAGKT<br>LDAQFENDERITPLESALMIWGSIEKEHDKLHEEIQNLIKIQAIAVCMENGNFKEAEEVFERIFGDPNSHMPFKSKLLMI<br>ISQKDTFHSFFQHFSYNHMMEKIKSYVNYVLSEKSSTFLMKAAAKVVESKSESAASSTGKEDKQPAPGPVEKPPREPARQ<br>LRNPPTTIGMMTLKAAFKTLSGAQDSEAAFAKLDQKDLVLPTQALPASPALKNKRPRKDENESSAPADGEGGSELQPKNK<br>RMTISRLVLEEDSQSTEPSAGLNSSQEAASAPPSKPTVLNQPLPGEKNPKVPKGKWNSSNGVEEKETWVEEDELFQVQAP<br>DEDSTTNITKKQKWTVEESEWVKAGVQKYGEGNWAAISKNYPFVNRTAVMIKDRWRTMKRLGMN |
| TRF2ΔTRFH | MGHHHHHHGTGSENLYFQGAGDSLEFIASKLAGSGSGSGMAGGGGSSDGSGRAAGRRASRSSGRARRGRHEPGLGGPAER<br>SESAASSTGKEDKQPAPGPVEKPPREPARQLRNPPTTIGMMTLKAAFKTLSGAQDSEAAFAKLDQKDLVLPTQALPASPA<br>LKNKRPRKDENESSAPADGEGGSELQPKNKRMTISRLVLEEDSQSTEPSAGLNSSQEAASAPPSKPTVLNQPLPGEKNPK<br>VPKGKWNSSNGVEEKETWVEEDELFQVQAPDEDSTTNITKKQKWTVEESEWVKAGVQKYGEGNWAAISKNYPFVNRTAVM<br>IKDRWRTMKRLGMN |
| TRF2cAcidic$_{TRF1}$ | MGHHHHHHGTGSENLYFQGAGDSLEFIASKLAGSGSGSGMAEDVSSAAPSPRGCADGRDADPTEEQMAETERNDEEQFEC<br>QELLECQVQVGPPEEEGAGEARLEEAVNRWVLKFYFHEALRAFRGSRYGDFRQIRDIMQALLVRPLGKEHTVSRLLRVMQ<br>CLSRIEEGENLDCSFDMEAELTPLESAINVLEMIKTEFTLTEAVVESSRKLVKEAAVIICIKNKEFEKASKILKKHMSKD<br>PTTQKLRNDLLNIIREKNLAHPVIQNFSYETFQQKMLRFLESHLDDAEPYLLTMAKKALKSESAASSTGKEDKQPAPGPV<br>EKPPREPARQLRNPPTTIGMMTLKAAFKTLSGAQDSEAAFAKLDQKDLVLPTQALPASPALKNKRPRKDENESSAPADGE<br>GGSELQPKNKRMTISRLVLEEDSQSTEPSAGLNSSQEAASAPPSKPTVLNQPLPGEKNPKVPKGKWNSSNGVEEKETWVE<br>EDELFQVQAPDEDSTTNITKKQKWTVEESEWVKAGVQKYGEGNWAAISKNYPFVNRTAVMIKDRWRTMKRLGMN |
| TRF2cHinge$_{TRF1}$ | MGHHHHHHGTGSENLYFQGAGDSLEFIASKLAGSGSGSGMAGGGGSSDGSGRAAGRRASRSSGRARRGRHEPGLGGPAER<br>GAGEARLEEAVNRWVLKFYFHEALRAFRGSRYGDFRQIRDIMQALLVRPLGKEHTVSRLLRVMQCLSRIEEGENLDCSFD<br>MEAELTPLESAINVLEMIKTEFTLTEAVVESSRKLVKEAAVIICIKNKEFEKASKILKKHMSKDPTTQKLRNDLLNIIRE<br>KNLAHPVIQNFSYETFQQKMLRFLESHLDDAEPYLLTMAKKALKRTRTITSQDKPSGNDVEMETEANLDTRKSVSDKQSA<br>VTESSEGTVSLLRSHKNLFLSKLQHGTQQQDLNKKERRVGTPQSTKKKKESRRATESRIPVSKSQPVTPEKHRAEDSTTN<br>ITKKQKWTVEESEWVKAGVQKYGEGNWAAISKNYPFVNRTAVMIKDRWRTMKRLGMN |
| TRF2cMyb$_{TRF1}$ | MGHHHHHHGTGSENLYFQGAGDSLEFIASKLAGSGSGSGMAGGGGSSDGSGRAAGRRASRSSGRARRGRHEPGLGGPAER<br>GAGEARLEEAVNRWVLKFYFHEALRAFRGSRYGDFRQIRDIMQALLVRPLGKEHTVSRLLRVMQCLSRIEEGENLDCSFD<br>MEAELTPLESAINVLEMIKTEFTLTEAVVESSRKLVKEAAVIICIKNKEFEKASKILKKHMSKDPTTQKLRNDLLNIIRE<br>KNLAHPVIQNFSYETFQQKMLRFLESHLDDAEPYLLTMAKKALKSESAASSTGKEDKQPAPGPVEKPPREPARQLRNPPT<br>TIGMMTLKAAFKTLSGAQDSEAAFAKLDQKDLVLPTQALPASPALKNKRPRKDENESSAPADGEGGSELQPKNKRMTISR<br>LVLEEDSQSTEPSAGLNSSQEAASAPPSKPTVLNQPLPGEKNPKVPKGKWNSSNGVEEKETWVEEDELFQVQAPDTPEKH<br>RARKRQAWLWEEDKNLRSGVRKYGEGNWSKILLHYKFNNRTSVMLKDRWRTMKKLKLISSDSED |
| TRF2ΔMyb | MGHHHHHHGTGSENLYFQGAGDSLEFIASKLAGSGSGSGMAGGGGSSDGSGRAAGRRASRSSGRARRGRHEPGLGGPAER<br>GAGEARLEEAVNRWVLKFYFHEALRAFRGSRYGDFRQIRDIMQALLVRPLGKEHTVSRLLRVMQCLSRIEEGENLDCSFD<br>MEAELTPLESAINVLEMIKTEFTLTEAVVESSRKLVKEAAVIICIKNKEFEKASKILKKHMSKDPTTQKLRNDLLNIIRE<br>KNLAHPVIQNFSYETFQQKMLRFLESHLDDAEPYLLTMAKKALKSESAASSTGKEDKQPAPGPVEKPPREPARQLRNPPT<br>TIGMMTLKAAFKTLSGAQDSEAAFAKLDQKDLVLPTQALPASPALKNKRPRKDENESSAPADGEGGSELQPKNKRMTISR<br>LVLEEDSQSTEPSAGLNSSQEAASAPPSKPTVLNQPLPGEKNPKVPKGKWNSSNGVEEKETWVEEDELFQVQAPD |
